# SRF-deficient astrocytes provide neuroprotection in mouse models of excitotoxicity and neurodegeneration

**DOI:** 10.1101/2023.05.17.541074

**Authors:** Surya Chandra Rao Thumu, Monika Jain, Sumitha Soman, Soumen Das, Vijaya Verma, Arnab Nandi, David H. Gutmann, Balaji Jayaprakash, Deepak Nair, James Premdoss Clement, Swananda Marathe, Narendrakumar Ramanan

## Abstract

Reactive astrogliosis is a common pathological hallmark of central nervous system (CNS) injury, infection, and neurodegeneration, where reactive astrocytes can be protective or detrimental to normal brain functions. Currently, the mechanisms regulating neuroprotective astrocytes and the extent of neuroprotection are poorly understood. Here, we report that conditional deletion of serum response factor (SRF) in adult astrocytes causes reactive-like hypertrophic astrocytes throughout the mouse brain. These *Srf*^GFAP-ER^CKO astrocytes do not affect neuron survival, synapse numbers, synaptic plasticity or learning and memory. However, the brains of *Srf* knockout mice exhibited neuroprotection against kainic-acid induced excitotoxic cell death. Relevant to human neurodegenerative diseases, *Srf*^GFAP-ER^CKO astrocytes abrogate nigral dopaminergic neuron death and reduce β-amyloid plaques in mouse models of Parkinson’s and Alzheimer’s disease, respectively. Taken together, these findings establish SRF as a key molecular switch for the generation of reactive astrocytes with neuroprotective functions that attenuate neuronal injury in the setting of neurodegenerative diseases.

## Introduction

CNS injury can be caused by diverse etiologies to result in either acute tissue damage (ischemia, traumatic brain injury) or chronic morphological and functional changes in neural tissues, resulting in behavioral and cognitive deficits as seen in neurodegenerative diseases (Burda and Sofroniew, 2014; Pekny and Pekna, 2014). Although much of the focus has centered on neuronal dysfunction, emerging studies have highlighted the critical roles played by non-neuronal cells, particularly astrocytes, in tissue repair, homeostasis, and disease progression (Burda et al., 2016; Kimelberg, 2010; Linnerbauer and Rothhammer, 2020; Pekny and Pekna, 2014). In this regard, astrocytes respond to CNS pathology by undergoing a spectrum of transcriptomal, physiological and structural changes, termed “reactive astrogliosis or reactive astrocytosis” (Burda and Sofroniew, 2014; Liddelow and Barres, 2017). Astrocyte reactivation is a diverse and complex cellular response that is context-dependent and reactive astrocytes perform several critical functions including aiding in repair and restoring normal homeostasis in the brain (Aswendt et al., 2022; Li et al., 2008; Linnerbauer and Rothhammer, 2020; Pekny and Pekna, 2014).

Although reactive astrocytes could provide neuroprotection in the initial stages of disease, prolonged gliosis could hamper normal neuronal functions and contributes to the pathophysiology of the disease (Gleichman and Carmichael, 2020; Huang et al., 2022; Pekny and Pekna, 2014; Phatnani and Maniatis, 2015; Sofroniew, 2014; Verkhratsky et al., 2016). For example, reactive astrocytes generated by lipopolysaccharide (LPS)-induced neuroinflammation are dependent on microglia, deficient in several critical astrocyte functions and cause death of neurons and oligodendrocytes (Guttenplan et al., 2021; Liddelow et al., 2017). Similarly, reactive astrocytes are also found in the aging brain and in the context of CNS neurodegeneration (Boisvert et al., 2018; Clarke et al., 2018; Liddelow et al., 2017), where inhibition of reactive astrocytes provides neuroprotection in mouse models of Parkinson’s disease, Alzheimer’s disease, and ALS (Ceyzeriat et al., 2018; Guttenplan et al., 2020b; Park et al., 2021; Reichenbach et al., 2019; Yun et al., 2018). For these reasons, suppression of reactive astrogliosis is actively being pursued as an astrocyte-targeted therapeutic strategy for the treatment of neurodegenerative diseases (Lee et al., 2022).

Previous studies have identified several genes including *Stat3*, *Fgfr*, *Aqp-4*, *Endothelin-1*, *β1-integrin* and *Bmal* whose deletion results in hypertrophic reactive-like astrocytes (Correa-Cerro and Mandell, 2007; Kang and Hebert, 2011; Lananna et al., 2018; Sofroniew, 2014). Furthermore, genetic manipulations of some of these genes have revealed their importance in regulating both the beneficial and negative effects of reactive astrocytes. For example, astrocyte-specific deletion of *Stat3* has revealed critical roles played by this pathway in the generation of scar-border forming astrocytes and tissue repair, and in the pathogenesis of Alzheimer’s and Huntington diseases (Abjean et al., 2023; Ben Haim et al., 2015; Herrmann et al., 2008; Okada et al., 2006; Reichenbach et al., 2019). Deletion of β1-integrin resulted in progressive astrogliosis and spontaneous seizures in adult mice (Robel et al., 2015; Robel et al., 2009). These observations raise the intriguing possibility that reactive astrocytes can be reprogrammed to improve neuronal survival and promote CNS repair and recovery following injury and in the setting of neurodegenerative diseases (Lee et al., 2022).

To identify potential mechanisms for reactive astrocyte reprogramming, we focused on SRF, a stimulus-dependent transcription factor that plays several critical roles in nervous system development and glial differentiation (Knoll et al., 2006; Knoll and Nordheim, 2009; Lu and Ramanan, 2011, 2012). We recently showed that astrocyte-specific deletion of SRF early during mouse development resulted in persistent reactive-like astrocytes throughout the postnatal mouse brain (Jain et al., 2021). Although these astrocytes did not cause any discernible abnormalities in the brain, the phenotypic changes exhibited by these *Srf*-deficient astrocytes could be due to developmental defects in astrocyte differentiation caused by SRF deletion (Jain et al., 2021; Lu and Ramanan, 2012). To address this, we now report that deletion of SRF in adult astrocytes also causes astrocyte reactivity-like phenotype that is persistent and widespread across the brain. We further show that *Srf-*deficient astrocytes did not affect neuronal survival, normal neuron functions, or learning and memory. Importantly, we demonstrate that astrocytic *Srf* deletion results in markedly attenuated neuronal death caused by excitotoxicity, and in a mouse model of Parkinson’s disease. Furthermore, astrocytic *Srf* deletion in the APP/PS1 mouse model of Alzheimer’s disease causes a significant decrease in β-amyloid plaque burden. Taken together, our results reveal SRF as a critical regulator of neuroprotective reactive astrogliosis in the context of brain injury and neurodegenerative disease.

## Results

### SRF deletion in adult astrocytes results in GFAP+ hypertrophic astrocytes

We recently showed that astrocyte-specific SRF deletion during embryonic development using a GFAP-Cre transgenic mouse line (*Srf*^GFAP^CKO) results in reactive-like astrocytes across the brain starting around 2 weeks of age and that these astrocytes persist throughout adulthood (Jain et al., 2021). These changes in astrocytes could reflect a developmental effect or result from indirect effects via other cell types with Cre-mediated *Srf* loss (e.g., neurons). To determine whether SRF functions in a cell-autonomous fashion in adult astrocytes to establish a non-reactive state, we generated *Srf*^f/f; GFAP-ERT+/-^ (*Srf*^GFAP-ER^CKO) mice in which the hGFAP promoter drives the expression of a tamoxifen-inducible Cre recombinase in postnatal astrocytes, rather than in GFAP-positive neural progenitor cells (fig. S1A) (Chow et al., 2008). In this hGFAP-Cre^ERT^ transgenic line, co-staining for β-gal from the Cre-IRES-β-gal transgene and cell-type specific marker genes revealed that ∼85% of GFAP+ and Sox9+ astrocytes were β-gal-positive in the neocortex, corpus callosum and hippocampus (fig. S1B, C, F). No Olig2+/β-gal+ oligodendrocyte lineage cells were found (fig. S1D, F), while fewer than 1% of NeuN+ cells were β-gal+ in the neocortex and striatum, and none were β-gal+ in the hippocampus (fig. S1E, F). To delete *Srf* in adult astrocytes, tamoxifen was administered to 6-8-week-old *Srf*^GFAP-ER^CKO mice and control littermates after astrocyte development was complete (Fig. 1A) (Wang and Bordey, 2008).

**Figure 1.**
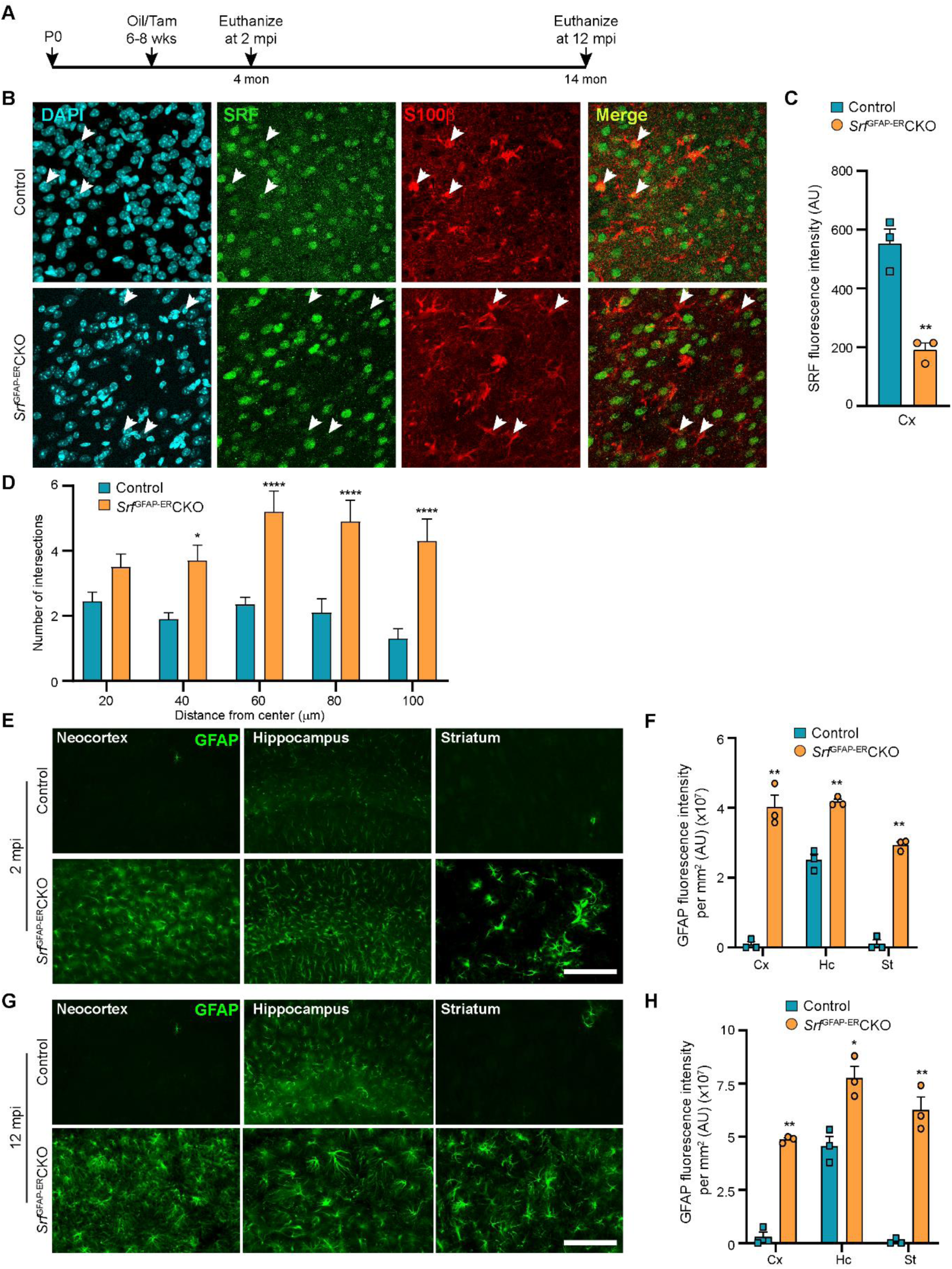
*Srf* deletion in post-natal astrocytes results in hypertrophic GFAP+ astrocytes. **(A)** Schematic of timeline of tamoxifen injection and analysis of astrocyte phenotype. **(B)** Coronal sections of neocortex showing immunostaining for SRF and S100β in control and *Srf*^GFAP-ER^CKO mice at 2 mpi. Astrocytes in control mice showed staining for both SRF and S100β while several astrocytes in the mutant mice did not show any SRF staining (white arrowheads). **(C)** Quantification of SRF fluorescence intensity in (B). **(D)** Sholl analysis of S100β expressing astrocytes from (B) shows hypertrophic morphology of SRF ablated astrocytes. n=3 mice. **(E)** Coronal sections showing immunostaining for GFAP in control and *Srf*^GFAP-ER^CKO mutant mice at 2 mpi. **(F)** Quantification of relative GFAP fluorescence intensity in F. n=3 mice. **(G)** Coronal sections showing immunostaining for GFAP in control and *Srf*^GFAP-ER^CKO mutant mice at 12 mpi. **(H)** Quantification of GFAP fluorescence intensity in H. n=3 mice. Data are represented as mean ± SEM. * P < 0.05, ** P < 0.01, **** P < 0.0001, ns, not significant. Unpaired t-test. Scale bar, 100 µm.

We first confirmed SRF deletion in the astrocytes in the *Srf*^GFAP-ER^CKO mice at 2 mpi by co-immunostaining for SRF and the astrocytic marker, S100β. The astrocytes in control mice showed robust expression of SRF while many astrocytes in the knockout mice did not show any SRF expression (Fig. 1B, C). We observed that the *Srf*-deficient astrocytes exhibited a hypertrophic morphology (Fig. 1D). Immunostaining for the astrocytic marker, GFAP, showed little to no expression in the neocortex and striatum and only basal expression in the hippocampal astrocytes of 2-month-old tamoxifen injected control mice. In contrast, the astrocytes in the tamoxifen-injected mutant mice exhibited robust GFAP expression in all the brain regions analyzed (Fig. 1E, F). Importantly, the associated phenotypic changes in astrocytes upon *Srf* deletion were not spatially restricted in the brain, as observed in studies using other knockout strains (Garcia et al., 2010; Kang et al., 2014). We next asked whether the increased GFAP expression and hypertrophic morphology of astrocytes was a transient phenomenon in the *Srf*^GFAP-ER^CKO mice or persisted throughout adulthood. While there were few to no GFAP-positive astrocytes in the neocortex and striatum, and weakly GFAP-positive astrocytes in the hippocampus of control mice at 12 mpi, intense GFAP-positive hypertrophic astrocytes were found in several regions of the brain, including the neocortex, hippocampus, and striatum of *Srf*^GFAP-ER^CKO mice (Fig. 1G, H). These findings reveal that *Srf* deletion in adult astrocytes also causes widespread and persistent GFAP+ hypertrophic astrocytes like that observed with embryonic deletion in astrocytes (Jain et al., 2021).

### *Srf*^GFAP-ER^CKO mice show upregulation of reactive astrocyte markers

Hypertrophic morphology with increased GFAP expression is generally exhibited by reactive astrocytes, which show a heterogeneous context-dependent transcriptomic profile reflective of their diverse reactive states (Das et al., 2020; Jiwaji et al., 2022; Zamanian et al., 2012). To study whether the *Srf*-deficient astrocytes exhibit reactivation, we performed immunostaining for the reactive astrocyte marker, vimentin along with GFAP (Ridet et al., 1997). We found that the GFAP-positive astrocytes in *Srf* cKO mice were also vimentin-positive, whereas no vimentin+/GFAP+ astrocytes were found in the control mice (Fig. 2A, B). We next analyzed the expression of genes that were described for reactive astrocytes induced in response to two acute pathological paradigms – lipopolysaccharide (LPS)-induced neuroinflammation and ischemic stroke (Liddelow et al., 2017; Zamanian et al., 2012). Expression levels were determined by real-time qRT-PCR of cortical tissue for the following genes: *Ugt1a1, LigP1, Serping1, Srgn, Psmb8, Fkbp5, Ggta1, Gbp2, Amigo2, Fbln5* (LPS group); and *Emp1, Clcf1, Slc10a6, CD109, CD14, Ptx4, S100a10, Ptgs2, B3gnt5, Tm4sf1, Sphk1, Tgm1* (ischemic stroke group); and *Gfap, Lcn2, Serpina3n, Aspg1, Cxcl10, Timp1* (pan-reactive markers). We observed increased expression of all pan-reactive markers tested in the *Srf*^GFAP-ER^CKO mouse brain compared to control mice (Fig. 2C). However, there were no discernible differences in the expression of genes that are upregulated in response to LPS versus ischemic stroke in the *Srf*^GFAP-ER^CKO mice. We found 6 of 12 genes in the LPS group and 9 of 12 genes in the stroke group were upregulated in the *Srf*^GFAP-ER^CKO mice (Fig. 2C). Together, this indicated that *Srf*-deficient astrocytes exhibit a heterogenous reactive-like phenotype.

**Figure 2.**
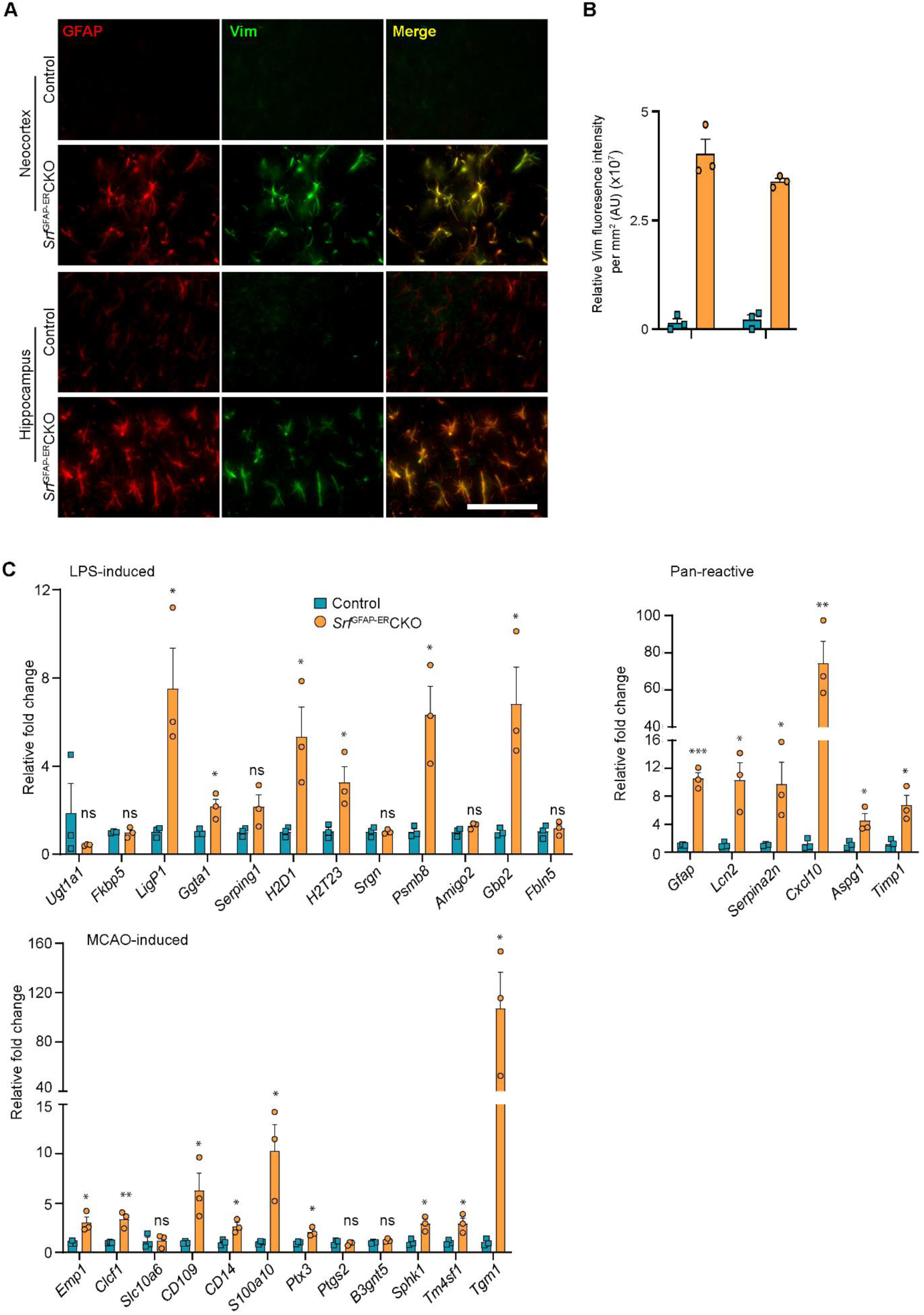
*Srf*-deficient astrocytes express reactive astrocyte markers. **(A)** Coronal sections showing GFAP and Vimentin (Vim) immunostaining in control and *Srf*^GFAP-ER^CKO mice at 2 mpi. Astrocytes in the neocortex and hippocampus were positive for both GFAP and Vimentin. The control astrocytes exhibited weaker staining for GFAP only in the hippocampus and not for Vimentin. **(B)** Quantification of Vim fluorescence intensity in the astrocytes in the neocortex and hippocampus shown in (A). n=3 mice. **(C)** Quantitative PCR analysis of genes induced by inflammatory LPS stimulation, middle cerebral artery occlusion (MCAO) and markers of pan-reactive astrocytes in control and *Srf* cKO mice. n=3 mice. ** P < 0.01. Unpaired t-test. Data are represented as mean ± SEM. Cx, neocortex; Hc, hippocampus. AU, arbitrary units. Scale bar, 50 µm.

### *Srf*-deficient astrocytes do not induce cell death in the brain

Since the *Srf*-deficient astrocytes showed expression of several marker genes associated with astrocyte reactivation, we next sought to determine any deleterious effects caused by *Srf*-deficient astrocytes. Brains sections from *Srf*^GFAP-ER^CKO mice and control littermates at 2 mpi and 12 mpi (4-months and 14-months of age, respectively) were stained using FluoroJade-C or TUNEL. We did not observe any discernible TUNEL- or FluoroJade-C-positive cells in the brains of *Srf*^GFAP-ER^CKO mice compared to control littermates (Suppl. Fig. S2A, B). To confirm the absence of cell loss, immunostaining for the neuronal marker, NeuN, showed no difference in the number of NeuN-positive cells in the neocortex, striatum, and hippocampus of *Srf*^GFAP-ER^CKO mice at 12 mpi (Fig. 3A, B). Similarly, immunostaining for the oligodendrocyte lineage marker, Olig2, revealed no difference in the number of Olig2^+^ cells in *Srf* mutant mice at 2 mpi relative to controls (Suppl. Fig. S3A, B). We next sought to determine whether this prolonged gliosis affected myelination in the brains of *Srf*^GFAP-ER^CKO mice. Black gold-II staining showed no discernible differences in myelin in the neocortex and hippocampus of *Srf*^GFAP-ER^CKO mice compared to control mice, even at 12 mpi (Suppl. Fig. S3C, D). Consistent with the lack of cell death, no gross morphological abnormalities were found in the brains of *Srf* cKO mice at 12 mpi (14-15 months of age) (Fig. 3A). Furthermore, the *Srf*^GFAP-ER^CKO mice did not exhibit any difference in body weight at 12 mpi compared to control littermates (Suppl. Fig. S3E). Taken together, these data demonstrate that SRF deletion in astrocytes does not cause neuronal cell death or affect myelination.

**Figure 3.**
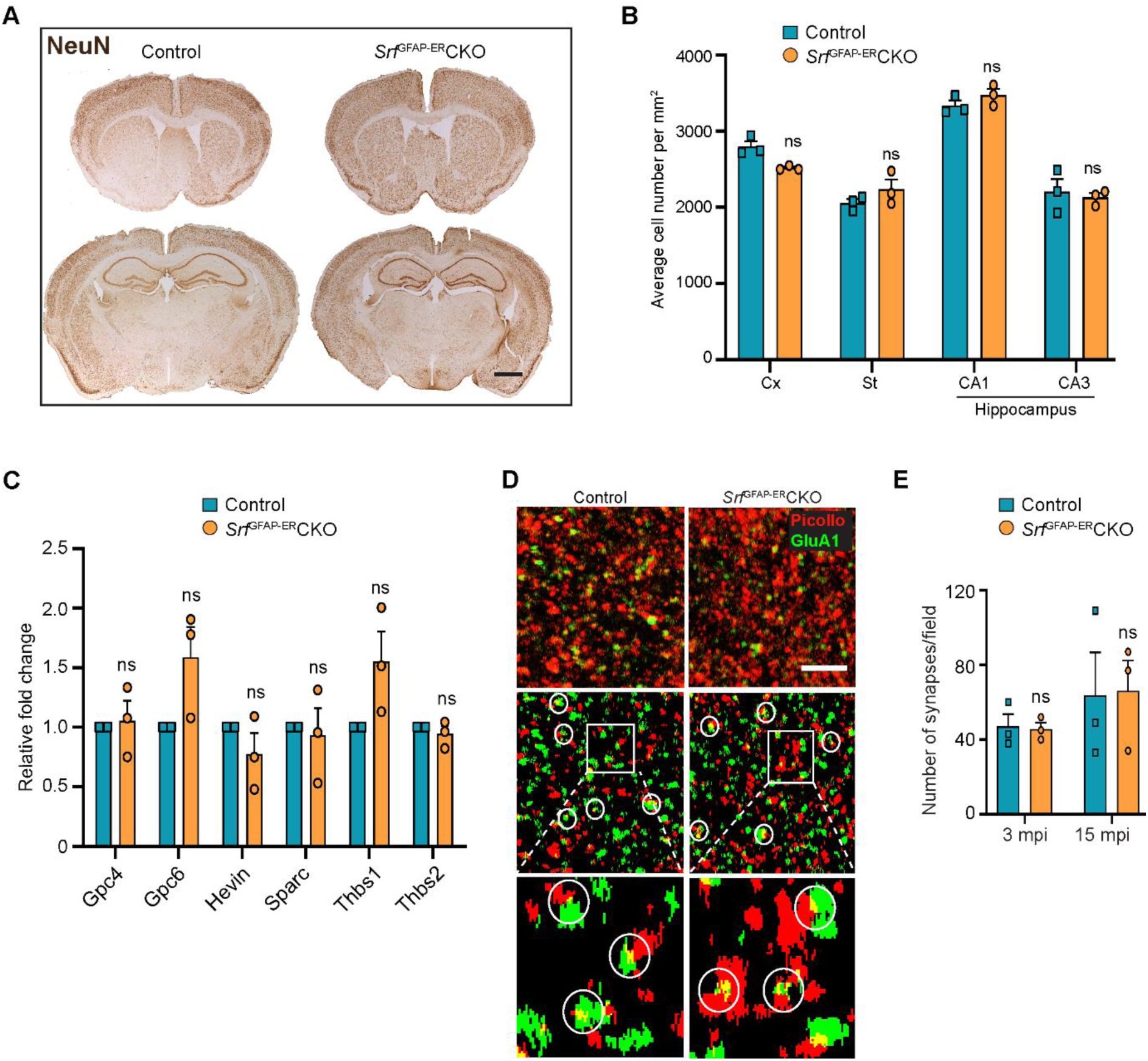
*Srf*-deficient astrocytes do not affect neuronal survival and synapse numbers. **(A)** Coronal sections showing NeuN immunostaining in control and *Srf* mutant mice at 12 mpi. **(B)** Quantification of NeuN+ cell numbers in the neocortex and striatum, and DAPI+ cell numbers in the CA1 and CA3 regions of the hippocampus. **(C)** Quantitative PCR of astrocyte-secreted synaptogenic factors shows no difference in their expression in the mutant mice when compared to control mice. n=3 mice. **(D)** Representative images of neocortical sections immunostained for the presynaptic marker, piccolo (red) and postsynaptic marker, GluA1 (green). Co-localization of staining (yellow puncta) was counted as a synapse. Scale bar, 10 µm. **(E)** Quantification of the number of synapses in the neocortex in control and *Srf*^GFAP-ER^CKO mutant mice at 3 and 15 mpi. n=3 mice. Unpaired t-test. Data are represented as mean ± SEM. ns, not significant.

Astrocytes play important roles in the maintenance of the blood-brain barrier (BBB) and abnormalities in their morphology or functions are associated with loss of BBB integrity (Abbott et al., 2006; Chapouly et al., 2015). Given the hypertrophic morphology and altered gene expression of astrocytes in the *Srf* cKO mice, we examined whether *Srf-*deficient astrocytes were compromised in their ability to support BBB integrity. Transcardial injection of a small 10 kDa dextran fluorescein tracer revealed no tracer in the brain parenchyma of *Srf*^GFAP-ER^CKO mice at 17-18-months of age (15 mpi) compared to control mice (Suppl. Fig. S4), suggesting no effect on BBB integrity.

Microglia rapidly respond to changes in the environment and there exists an active crosstalk between astrocytes and microglia (Jha et al., 2019; Matejuk and Ransohoff, 2020). Given the morphological and molecular changes exhibited by *Srf*-deficient astrocytes, we next examined the status of microglia in the *Srf*^GFAP-ER^CKO mice. In the healthy brain, Iba1 immunostaining shows basal expression in the microglial cell body and in highly ramified processes, indicative of a resting state. We observed a similar pattern of immunostaining, morphology, and fluorescence intensity in *Srf* cKO mouse brains at 2 mpi (Suppl. Fig. S5A, B). Cell counts further revealed similar numbers of Iba1-positive microglia in the neocortex and hippocampus of *Srf* cKO mice at 2 mpi relative to control littermates (Suppl. Fig. S5A, B). However, Iba1 immunostaining at 12 mpi revealed an increase in fluorescence intensity accompanied with morphological changes in *Srf* cKO mice, including thickening of the processes and the acquisition of an amoeboid shape, compared to control mice (Suppl. Fig. S5C, D). Cell counts at 12 mpi showed an increase in IbaI-positive cells per unit area in the neocortex, hippocampus, and striatum of *Srf* cKO mice relative to control mice (Suppl. Fig. S5C, D). Moreover, neither astrocytes nor microglia were actively proliferating (phospho-histone H3 (phH3) immunoreactivity) (Suppl. Fig. S2C) and the increased microglial cell numbers in the *Srf* cKO mice at 12 mpi likely occurred at an earlier time point.

### *Srf*^GFAP-ER^CKO mice do not exhibit deficits in synaptic plasticity and behavior

Astrocytes play major roles in synapse formation, maintenance, and elimination (Augusto-Oliveira et al., 2020; Baldwin and Eroglu, 2017; Chung et al., 2015). We therefore asked whether *Srf*-deficient astrocytes are compromised in their ability to support synapse formation and function. First, we assessed mRNA expression of several astrocyte-secreted synaptogenic factors (Allen and Eroglu, 2017). Quantitative real-time PCR analyses showed no change in the expression levels of *Hevin, Glycipan 4/6, Thbs1,* and *Thbs2*, suggesting that the *Srf*-deficient astrocytes are not compromised in their ability to make pro-synaptogenic factors (Fig. 3C). Second, we investigated whether synapse numbers were altered in the *Srf*^GFAP-ER^CKO mice. Brain slices from 3 mpi and 15 mpi control and *Srf*^GFAP-ER^CKO mice were co-labeled with Piccolo (presynaptic marker) and GluA1 (post-synaptic marker) antibodies, and the number of synapses quantified based on overlapping puncta staining (Harris and Weinberg, 2012). Similarly, there was no difference in the number of synapses in *Srf*^GFAP-ER^CKO mice relative to control littermates (Fig. 3D, E), arguing that synapses are not affected by *Srf*-deficient astrocytes. Third, we asked whether synaptic transmission or synaptic plasticity are affected in *Srf*^GFAP-ER^CKO mice. Basal synaptic transmission, paired-pulse ratio, and long-term potentiation (LTP) were measured in hippocampal slices (Schaffer-collateral pathway) obtained from *Srf*^GFAP-ER^CKO and control littermates at 3 mpi and 15 mpi (Booth et al., 2014; Zaman et al., 2000). We did not observe any significant differences in either basal synaptic transmission or LTP in the knockout mice compared to their control littermates (3 mpi: synaptic transmission, p=0.3698; LTP, p value=0.5306; 15 mpi: synaptic transmission, p=0.1411; LTP, p value=0. 07334) (Fig. 4A-D and Suppl. Fig. S6). Taken together, these findings indicate that *Srf*-deficient astrocytes do not affect synaptic transmission or synaptic plasticity.

**Figure 4.**
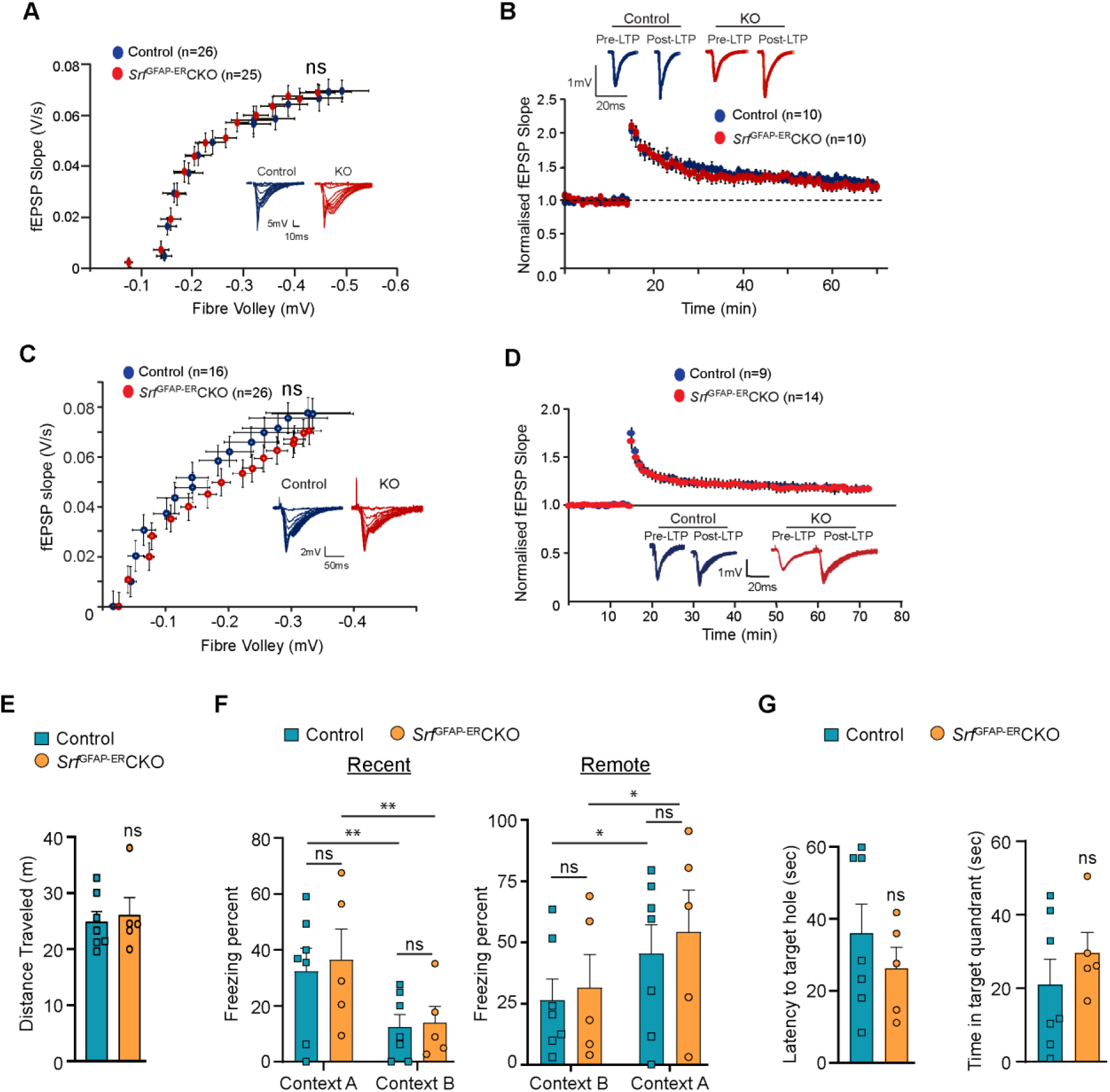
*Srf*-deficient astrocytes support normal synaptic plasticity, learning and memory. **(A-D)** Electrophysiological measurements showed no significance between control and mutant mice. **(A)** Basal synaptic transmission, **(B)** LTP in the hippocampus at 3 mpi, **(C)** Basal synaptic transmission, **(D)** LTP in the hippocampus at 15 mpi. Example traces are those recorded for 1-2 minutes around the time point indicated by I and II in the graph. The number of slices recorded from are indicated in parentheses. **(E-G)** Behavioral analyses. **(E)** Open-field test, **(F)** contextual fear conditioning at recent and remote time points in context A (shock context) and context B (no-shock context), and **(G)** Barnes maze test at 9 mpi in control and mutant mice. n=7 (control), n=5 (mutant) mice. Data are represented as mean ± SEM. ns, not significant. Unpaired t-test; 2-way ANOVA, Sidak’s post hoc test (F).

We next examined the effect of SRF deficiency on hippocampus-dependent spatial memory. To control for the confounding effects of possible changes in locomotor behavior, we performed an open field test and found no significant differences in the total distance traveled between *Srf*^GFAP-ER^CKO mice and the control littermates (control, 24.83 ± 1.85 m; *Srf*^GFAP-ER^CKO, 26.05 ± 3.10 m; p=0.74, unpaired t-test) (Fig. 4E). Next, we used the contextual fear-conditioning paradigm to assess spatial memory in *Srf*^GFAP-ER^CKO mice and control littermates at 9 mpi. 24 hours following fear conditioning in a shock context (context A), animals were tested for their freezing response in the same context, followed by context B which served as a control environment which wasn’t paired with the foot shock. While we found a statistically significant difference in the percent time spent freezing between context A and context B across genotypes (F (1, 10) = 26.42, p=0.0004), we did not find any difference between the genotypes in their freezing responses (Context A: control, 32.44 ± 8.20; *Srf*^GFAP-ER^CKO, 36.53 ± 10.99, p=0.91, Context B: control, 12.51 ± 4.37; *Srf*^GFAP-ER^CKO, 13.87 ± 5.91, p=0.99, 2-way ANOVA, Sidak’s *post hoc* test) (Fig. 4F). The mice were tested again in the same contexts (context B and context A) for remote recall after 1 month from fear conditioning. Again, while both genotypes could differentiate between the 2 contexts (F (1, 10) = 15.73, p=0.0027), we found no difference in the percent time spent freezing between *Srf*^GFAP-ER^CKO mice and control littermates in either of the contexts (Context B: control, 26.41 ± 8.55; *Srf*^GFAP-ER^CKO, 31.64 ± 13.42, p=0.95; Context A: control, 45.40 ± 11.86, *Srf*^GFAP-ER^CKO, 54.31 ± 17.08, p=0.86, 2-way ANOVA, Sidak’s *post hoc* test) (Fig. 4F).

Finally, we assessed spatial memory in the Barnes maze test. Animals were trained to locate an escape box on a Barnes maze until the learning curve flattened. Mice were tested in a probe test in the absence of an escape box, 2 hours after the last trial, and the latency to locate the target hole and the time spent in the target quadrant were calculated. We did not find a significant difference in the latency to target hole location (control, 35.95 ± 8.08, *Srf*^GFAP-ER^CKO: 26.22 ± 5.90, p=0.35, unpaired t-test) and the time spent in the target quadrant between *Srf*^GFAP-ER^CKO mice and control littermates (control, 20.96 ± 6.90; *Srf*^GFAP-ER^CKO, 29.60 ± 5.64, p=0.35, unpaired t-test) (Fig. 4G). Collectively, these experiments establish that the persistent and widespread presence of *Srf*-deficient astrocytes is not detrimental to normal synaptic plasticity or learning and memory.

### Transcriptomic profile of *Srf*-deficient astrocytes

To gain a better understanding of the molecular nature of *Srf*-deficient astrocytes, we performed RNA sequencing (RNA-seq) analysis of astrocytes isolated from the forebrain of 5-week-old *Srf*^GFAP^CKO (Jain et al., 2021) and 4 mpi (6-month-old) *Srf*^GFAP-ER^CKO mice and their respective control littermates. Genes with fold change of > 1.5 and adjusted p-value < 0.05 were considered significant. The expression profile of differentially expressed genes (DEGs) across the samples is presented in heatmaps (Fig. 5A). There were 315 common up-regulated genes in the *Srf*^GFAP^CKO and *Srf*^GFAP-ER^CKO astrocytes. Enrichment analysis of Gene Ontology terms for Biological Process (GO BP) indicated that SRF deficiency results in enrichment of pathways related to immune response and innate immunity (*Cd86, H2-T23, Cd84, Lst1, Ifit3, Trim30a, Trim30b*), inflammatory response (*Nlrp1b, Ccl12, Cxcl13, Ccl5, Ccl2, Cd14*), antiviral defense (*Zbp1, Rsad2, Mx1, Oas1a, Oas1g, Oas2*), response to interferon-beta and regulation of interleukin-1 beta (Fig. 5B, D; Suppl. Fig. S7A). There was also an enrichment of genes related to microglial cell activation (*Tlr1, C1qa. Itgam, Tlr6, Tlr2*) (Fig. 5B, D; Suppl. Fig. S7A).

**Figure 5.**
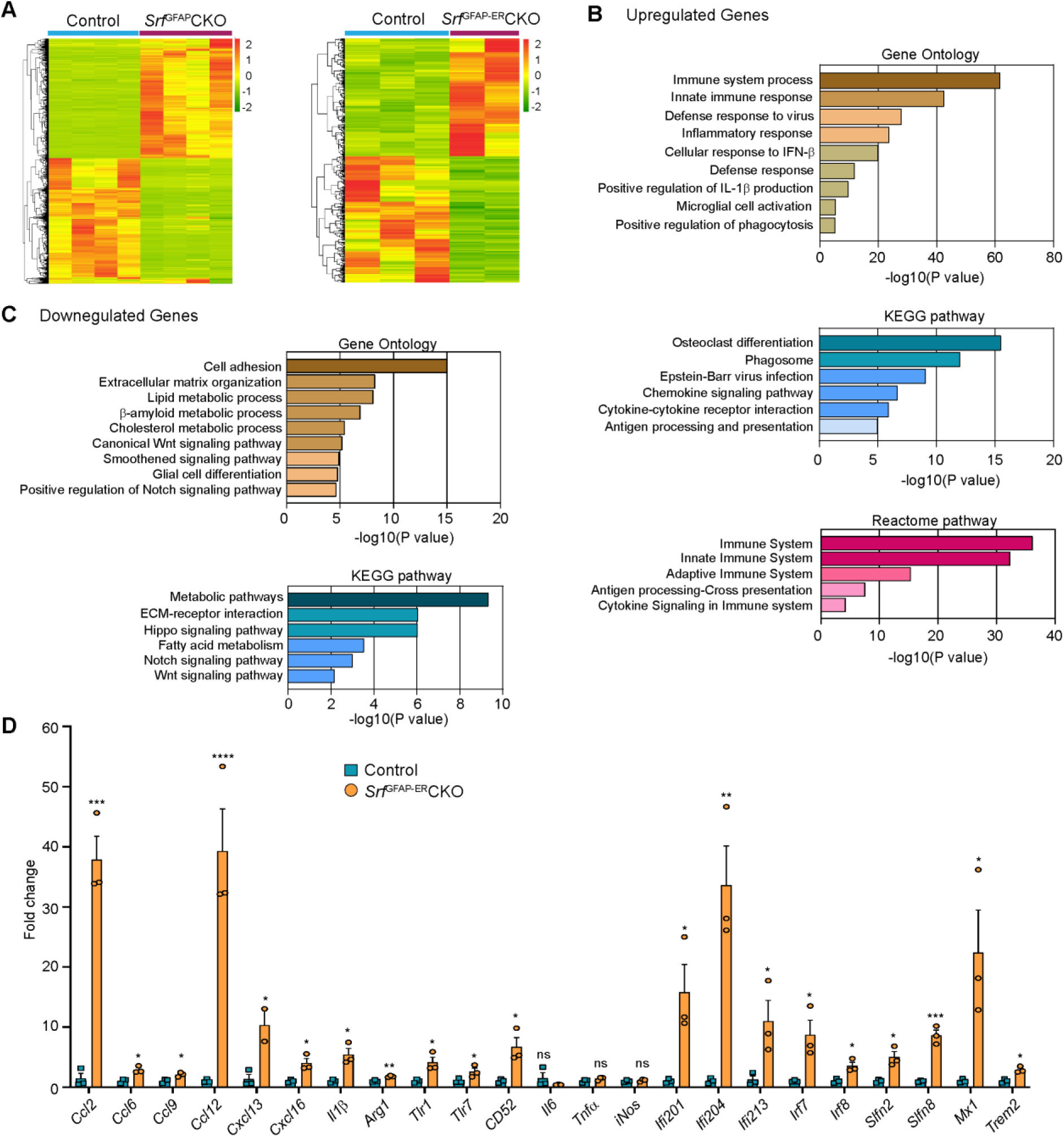
Transcriptome profile of *Srf*-deficient astrocytes. **(A)** Heat-maps of differentially expressed genes in control and SRF-deficient astrocytes from RNA sequencing. **(B)** Gene Ontology, KEGG pathway and Reactome pathway analysis of upregulated genes in *Srf*-deficient astrocytes. **(C)** Gene Ontology and KEGG pathway analysis of downregulated genes in *Srf*-deficient astrocytes. **(D)** Quantitative PCR analysis of interferon, inflammatory and defense response genes in reactive astrocytes identified in Gene Ontology analysis. n=3 mice. Data are represented as mean ± SEM. * P < 0.05, ** P < 0.01, *** P < 0.005, ****P < 0.001, ns, not significant. Unpaired t-test.

Analysis of downregulated genes revealed pathways related to cell adhesion (*Pcdhgb5, lamc1, Hapln3, Celsr2*), extracellular matrix (*Ecm2, Eln, Dmp1, Fnln1*), lipid metabolism (*Slc27a1, Acaa2, Msmo1, Aacs, FADS2*), Wnt signaling pathway (*Yap1, Fzd1, Fzd2, Wnt7b*), smoothened signaling pathway (*Evc2, Ptch1, Tulp3, Pax6, Gpr37l1, Gli1, Gli3, Gli2*) and Notch signaling pathway (*Sox2, Dll4, Enho, Yap1, Notch1, Hes1, Ascl1, Hes5*) (Fig. 5C, Suppl. Fig. S7B). Genes involved in glial cell differentiation (*Notch1, Chd2, Erbb2, Ascl1, Klf15*) were also downregulated, consistent with our earlier observations on the role of SRF in regulating glial differentiation (Lu and Ramanan, 2012). Under certain conditions, reactive astrocytes have been shown to cause cell death of neurons and oligodendrocytes partly through secretion of neurotoxic factors such as lipocalin 2 (*Lcn2*) and saturated fatty acids (Bi et al., 2013; Guttenplan et al., 2021). We analyzed our expression data set and found that the expression of Lcn2, C3 and the elongase of very-long fatty acid (Elovl) family of genes (involved in the synthesis of longer-chain saturated fatty acids) either remained unchanged or were downregulated in *Srf*-deficient astrocytes (Suppl. Fig. S7C). Together, these data suggested that *Srf*-deficient astrocytes exhibit a distinct transcriptome, and do not exhibit upregulation of genes shown to be involved in neurotoxicity.

### *Srf*-deficient astrocytes show up-regulation expression of genes involved in neuroprotection

Next, we analyzed the RNA sequencing data for potential genes that could provide insights into the potential functional nature of these astrocytes. The transmembrane chemokine CXCL16 has been shown to promote physiological neuroprotection following ischemic and excitotoxic insults via astrocytic release of CCL2 and adenosine 3 receptor (Adora3/A3R) (Rosito et al., 2012; Rosito et al., 2014). Expression of *Cxcl16, Ccl2* and *Adora3/A3R* expression was increased in *Srf*-deficient astrocytes (log_2_fold change: *Cxcl16*, 4.3-fold; *Ccl2,* 4.1-fold; *Adora3,* 3.3-fold) (Fig. 5D; Suppl. Fig. S8A, B). Insulin-like growth factor1 (IGF-1) was also increased 3.2-fold, which has been shown to provide neuroprotection in models of kainic-acid induced excitotoxicity (Chen et al., 2019). We also found an upregulation of genes involved in oxidative defense system, namely glutathione peroxidase-1 (GPx1) (2-fold) and heme oxygenase 1 (*Hmox1/HO-1*) (1.8-fold) (Suppl. Fig. S8C, D). Enrichment analysis of Gene Ontology terms for Biological Process (GO BP) revealed that *Srf*-deficient astrocytes showed enrichment of pathways related to cellular response to beta-amyloid and beta-amyloid clearance. Among the cellular response to beta-amyloid genes, *NAMPT* (2.9-fold)*, Casp4* (4.6-fold)*, Trem2* (5.5-fold)*, IGF-1* (3.2-fold) and *Adrb2* (2.4-fold) were upregulated. In addition, genes that promote Aβ uptake and clearance were also increased. These include lipoprotein lipase (*Lpl*) (4-fold), *HO1/Hmox-1* (1.8-fold), the member of the superfamily of ATP-binding cassette (ABC) transporters *Abca1* (1.7-fold) and beta-2-adrenergic receptor (*Adrb2*) (2.4-fold) and IGF-1 (Suppl. Fig. S8E, F). Also, genes whose downregulation has been shown promote Aβ production or clearance, *Bace-2* (−3.5-fold) and *Srebf2* (−2.1-fold) were downregulated in *Srf*-deficient astrocytes (Suppl. Fig. S8E, F).

### *Srf*^GFAP-ER^CKO mice exhibit reduced excitotoxic cell death

We finally sought to determine whether *Srf*-deficient astrocytes could provide neuroprotection in response to neuronal insult or in the setting of disease. Neuronal excitotoxicity is a major cause of cell death in CNS injuries and neurodegenerative diseases (Binvignat and Olloquequi, 2020). Kainic acid (KA) causes neurotoxic cell death of mainly the hippocampal CA1 and CA3 pyramidal neurons (Pollard et al., 1994). We asked whether SRF-deficient astrocytes could protect neurons from KA-induced excitotoxicity. We administered KA via intracerebroventricular injection into the lateral ventricles of control and mutant mice at 4 mpi (Fig. 6A) and assessed cell death using TUNEL and FluoroJade-C staining at 7 days post-KA injection. While control mice showed numerous TUNEL+ and FluoroJade-C+ cells in the CA1, CA3 and dentate gyrus, indicative of extensive neuronal cell death (Fig. 6B, C and Suppl. Fig. S9A-C), little to no cell death was seen in the hippocampus of *Srf*^GFAP-ER^CKO mice (Fig. 6B, C and Suppl. Fig. S9A-C).

**Figure 6.**
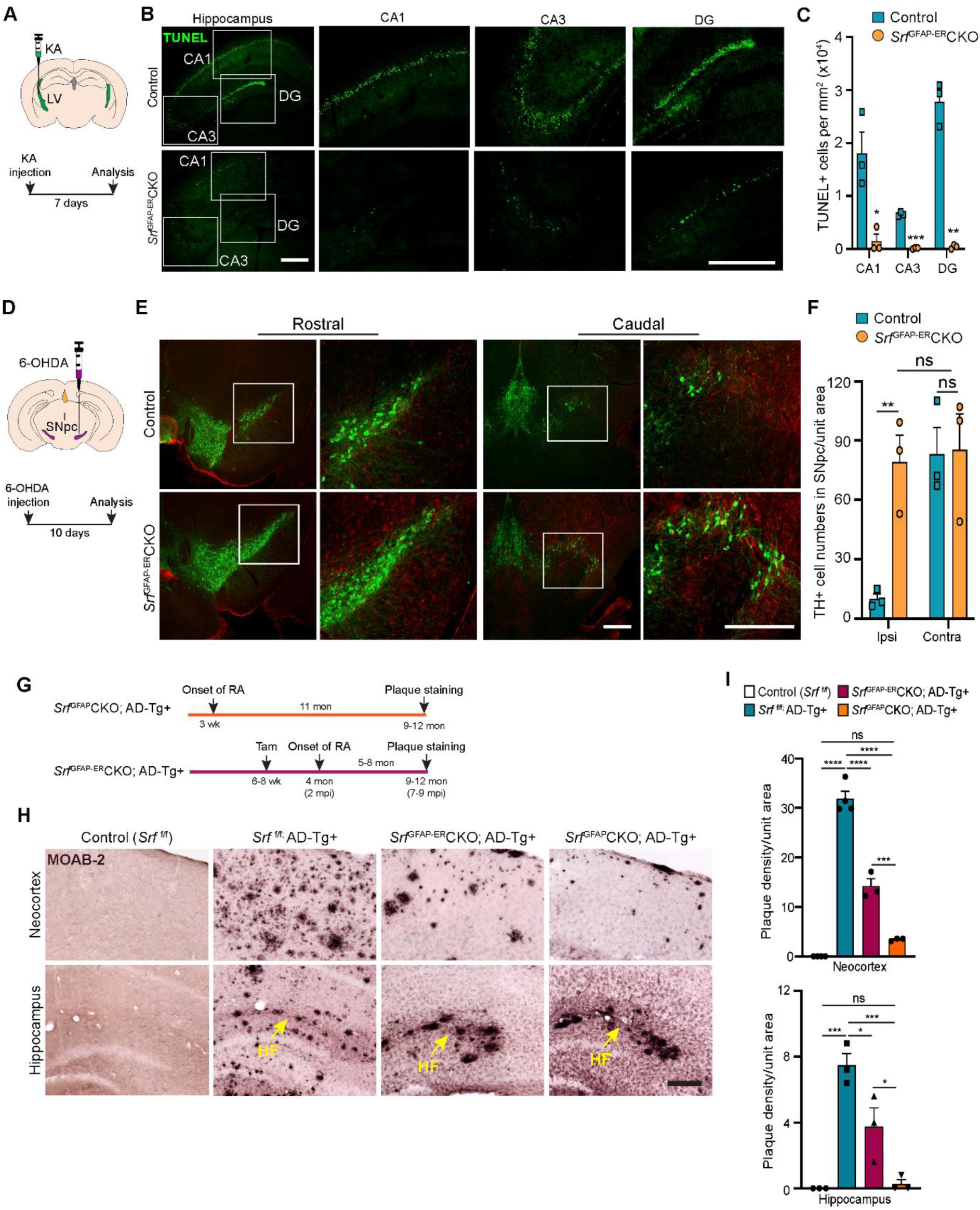
*Srf*-deficient reactive astrocytes confer neuroprotection in the brain. **(A)** Schematic of stereotaxic intracerebroventricular (ICV) injection of kainic acid (KA) (top) and timeline of experiment (bottom) in 4 mpi mice. **(B)** Coronal brain sections showing TUNEL (terminal deoxynucleotidyl transferase dUTP nick end labeling) staining at 7 days post-kainic acid injection shows cell death in hippocampal regions CA1, CA3 and DG. Scale bar, 200 µm. **(C)** Quantification of TUNEL^+^ cells in the hippocampal regions in B. n=4 mice. **(D)** Schematic of stereotaxic injection 6-hydroxydopamine (6-OHDA) in the substantia nigra *pars compacta* (SNpc) (top) and timeline of experiment (bottom) in control and mutant mice at 9 mpi. **(E)** Immunostaining of coronal sections showing tyrosine hydroxylase (TH^+^) dopamine neurons and GFAP^+^ astrocytes in the rostral and caudal regions of substantia nigra at 10 days following 6-OHDA injection. Magnified image of the white boxed region is shown on the right. Scale bar, 200 µm. **(F)** Quantification of TH^+^ neurons in the ipsilateral (ipsi) and contralateral (contra) side of (E). **(G)** Schematic showing the timeline of reactive astrocyte (RA) induction and amyloid plaque staining in AD mice. **(H)** Coronal sections showing MOAB2-immunostained β-amyloid plaques. Scale bar, 200 µm. **(I)** Quantification of plaque density from (H). RA, reactive astrocytes, LV, lateral ventricular, SNpc, Substantia nigra pars compacta, SR, Stratum radiatum. Yellow arrows show hippocampal fissure, HF. n=3 mice. Data are represented as mean ± SEM. * P < 0.05, ** P < 0.01, *** P < 0.005, **** P < 0.001, ns, not significant. Unpaired t-test (C, F), One-way ANOVA with Tukey *post-hoc* test (I).

### *Srf*-deficient astrocytes protect dopaminergic neurons in a model of Parkinson’s disease

Parkinson’s disease (PD) is caused by death of midbrain dopaminergic (DA) neurons in the substantia nigra pars compacta region (SNpc). We employed the 6-hydroxydopamine (6-OHDA) model of PD to investigate whether *Srf*-deficient reactive astrocytes can protect DA neurons from cell death (Ungerstedt, 1968). We first confirmed the presence of reactive astrocytes in the substantia nigra region in the *Srf* cKO mice before administration of 6-OHDA (Suppl. Fig. S10A, B). Next, unilateral stereotaxic injection of 6-OHDA was administered into the SNpc of control and mutant mice at 9 mpi and cell death was analyzed at 10 days post-injection for cell death (Fig. 6D). Control mice showed significant loss of tyrosine hydroxylase (TH)-positive DA neurons in the ipsilateral side compared to uninjected contralateral size (Fig. 6E, F and Suppl. Fig. S10C, D). In striking contrast, there was little to no DA neuron and fiber loss in the ipsilateral side of *Srf*^GFAP-ER^CKO mice compared to the ipsilateral side of control and to the contralateral side of knockout mice (Fig. 6E, F and Suppl. Fig. S10C, D).

### *Srf*^GFAP-ER^CKO/AD transgenic mice exhibit reduced amyloid plaque burden

We next investigated the role of reactive astrogliosis in the APP_Swe_/PSen1_dE9_ (APP/PS1) model of Alzheimer’s disease (AD). A morphological hallmark of AD is the deposition of β-amyloid (Aβ) plaques, which is associated with neurodegeneration and cognitive decline. We intercrossed *Srf*^GFAP-ER^CKO mice and APP/PS1 transgenic mice to generate *Srf*^GFAP-ER^CKO; AD^Tg+/-^ triple transgenic mice. *Srf*^GFAP-ER^CKO; AD^Tg+/-^ mice and control mice received tamoxifen injections at 6-8 weeks of age, and all groups of mice were analyzed at 9-12 months of age for Aβ plaques (Fig. 6G). Immunostaining using the MOAB-2 antibody, which detects only amyloid-β peptide, but not APP (Youmans et al., 2012), showed numerous plaques in the neocortex and hippocampus of APP/PS1 control mice (*Srf*^f/f^; AD^Tg+/-^) (Fig. 6H, I). In contrast, *Srf*^GFAP-ER^CKO; AD^Tg+/-^ mice had 60% fewer plaques compared to APP/PS1 mice (Fig. 6H, I). In *Srf*^GFAP-ER^CKO; AD^Tg+/-^ mice, reactive astrocytes were present for 5-8 months (until the end of the experiment) starting at 4 months of age. We then sought to determine whether *Srf*-deficient astrocytes, if present for a longer duration, can cause further reduction in the number of plaques. To address this, we crossed the *Srf*^GFAP^CKO mice with APP/PS1 mice to generate *Srf*^GFAP^CKO; AD^Tg+/-^ transgenic mice. In *Srf*^GFAP^CKO mice, SRF deletion occurs embryonically (around E16.5), and reactive-like astrocytes are seen starting at 3 weeks of life and persist throughout adulthood (Jain et al., 2021). Surprisingly, *Srf*^GFAP^CKO; AD^Tg+/-^ mice greatly reduced number of plaques in the neocortex and hippocampus (Fig. 6H, I), demonstrating that *Srf*-deficient astrocytes aid in efficient plaque reduction and that longer presence of these mutant astrocytes promoted greater reduction in amyloid plaques.

## Discussion

Astrocyte reactivity is a critical cellular response that plays a pivotal role in aiding neuronal recovery and restoring normal functions following injury and in neurodegenerative diseases (Pekny and Pekna, 2014; Sofroniew, 2005). Whether reactive astrocytes generated in response to tissue damage and in the setting of disease are beneficial or detrimental for normal brain health is context-dependent (Moulson et al., 2021; Pekny and Pekna, 2014; Sofroniew, 2020). Mounting evidence has shown that reactive astrocytes often become detrimental to normal neuronal functions and can act as active drivers of neurodegeneration (Huang et al., 2022; Phatnani and Maniatis, 2015). Therefore, it is critical to identify factors that can modulate astrocyte reactive states to achieve beneficial outcomes and provide therapeutic targets to treat neuronal injuries and neurodegenerative diseases. In this study, we demonstrate that conditional SRF ablation in adult astrocytes causes widespread reactive-like astrocytes in the brain. We show that SRF-deficient astrocytes do not affect neuronal and oligodendrocyte survival, BBB integrity, synapse numbers, synaptic plasticity, or learning and memory. Strikingly, these *Srf*-deficient reactive astrocytes protect hippocampal neurons from excitotoxicity and nigral dopaminergic neurons from 6-OHDA-induced cell death, as well as reduce amyloid plaque burden in a mouse model of AD.

SRF is a ubiquitous transcription factor that plays several critical functions in the nervous system (Tabuchi and Ihara, 2022). SRF ablation in neurons caused deficits in cell migration, neuronal structure and connectivity, synaptic plasticity and learning and memory (Johnson et al., 2011; Knoll and Nordheim, 2009; Lu and Ramanan, 2011). SRF deletion in developing neural stem cells specifically affected glial differentiation (Lu and Ramanan, 2012). Our observations show that SRF deletion in perinatal or adult astrocytes causes astrocyte reactivity-like state in the brain suggesting that SRF plays an important non-cell autonomous role in maintaining astrocytes in a non-reactive state (Jain et al., 2021). Other genes have been reported to regulate astrocyte nonreactive state albeit in a region-restricted manner, but whether these mutant astrocytes persist in a reactive-like state throughout adulthood is not known (Garcia et al., 2010; Kang et al., 2014).

Reactive astrocytes, depending on the context of injury and neurodegeneration, exhibit a spectrum of phenotypic changes that could alleviate or exacerbate neuropathology (Moulson et al., 2021). Reactive astrocytes observed in neuroinflammatory conditions and neurodegenerative disorders are neurotoxic and have degenerative influences on the brain (Boisvert et al., 2018; Chun et al., 2020; Clarke et al., 2018; Liddelow et al., 2017; Yun et al., 2018). In fact, inhibition of the formation of these neuroinflammatory reactive astrocytes has been found to be beneficial for neuronal survival following injury and attenuated neurodegeneration in mouse models of Alzheimer’s disease, Parkinson disease and ALS (Guttenplan et al., 2020a; Guttenplan et al., 2020b; Mann et al., 2022; Park et al., 2021; Yun et al., 2018). On the other hand, reactive astrogliosis following ischemic stroke and traumatic brain injury exhibits neuroprotective functions (Aswendt et al., 2022). Our findings show that *Srf*-deficient astrocytes are neuroprotective in the brain. These mutant astrocytes efficiently protected hippocampal neurons from excitotoxicity. The cell populations that are most vulnerable to systemic kainic acid-induced cell death are the hippocampal CA3 and CA1 pyramidal neurons while the dentate granule neurons are usually spared (Sperk, 1994). We found that intracerebroventricular injection of kainic acid caused death of neurons in all the regions including the dentate gyrus in the control mice and this was attenuated in the *Srf* cKO mice. Death of CA3 neurons is due to excitotoxicity while that of CA1 neurons is thought to be due to anoxia/ischemia caused by status epilepticus (Sperk, 1994). The decreased cell death in these regions in the *Srf* cKO mice suggests that *Srf*-deficient astrocytes likely protect neurons from both anoxic/ischemic and excitotoxic cell death. 6-OHDA is an oxidizable analog of dopamine that causes death of nigral dopaminergic (DA) neurons via oxidative stress (Hernandez-Baltazar et al., 2017). The lack of any discernible cell death of DA neurons following 6-OHDA administration in the *Srf* cKO mice revealed robust neuroprotection from oxidative stress. In addition, *Srf*-deficient astrocytes in the APP/PS1 model of AD resulted in a greater reduction of amyloid plaque burden in the cortex and hippocampus. These observations together suggest that SRF is a major mediator of reactive astrocytes that could provide neuroprotection in the brain following excitotoxity and in neurodegeneration.

There are some limitations to the current study. We have shown that the *Srf* cKO mice do not exhibit any gross structural abnormalities or deficits in synaptic plasticity and behavior. However, it is possible that they may exhibit as yet uncharacterized structural or behavioral abnormalities. Astrocyte-restricted β1-integrin deletion results in widespread reactive astrocytes but these mutant mice showed spontaneous seizures (Robel et al., 2015). Although there was no obvious occurrence of seizures in the *Srf* cKO mice, electroencephalography recordings may provide information on any neuronal hyperexcitability in these mutant mice. Another limitation is that the molecular mechanisms by which *Srf*-deficient astrocytes provide neuroprotection is not known. The *Srf*-deficient astrocytes upregulate genes involved in regulating oxidative stress and these, including *Cxcl16-Ccl2-Adora3*, along with *GPx1, Hmox-1* could protect neurons from kainate-induced excitotoxicity and following 6-OHDA administration. We also observed a significant reduction in Aβ plaque burden, and this could be due to either deficits in Aβ production or efficient clearance by astrocytes. Several genes involved in Aβ clearance were upregulated in *Srf*-deficient astrocytes suggesting that astrocytes may promote clearance. We observed a greater accumulation of plaques near the cortical pial surface and in the hippocampal fissure region and these could be potential regions through which Aβ is cleared. Microglia can also become reactive in response to injury and neurodegeneration and reactive microglia can exert beneficial or detrimental effects on neuronal repair and recovery (Muzio et al., 2021). The molecular or functional nature of the reactive microglia observed in the *Srf* cKO mouse brains is not known. The absence of any structural and behavioral abnormalities in the *Srf*^GFAP-ER^CKO mice strongly suggests that altered microglial morphology and increased numbers in these mice are unlikely to be detrimental and could potentially contribute to neuroprotection and Aβ clearance.

In summary, using the SRF transcription factor as a key regulator of reactive astrocytes, we demonstrate that the persistent presence of *Srf*-deficient astrocytes is not detrimental to brain cell survival, architecture, synaptic plasticity, learning or memory. Importantly, in the setting of both excitotoxicity and neurodegenerative disease these *Srf*-deficient reactive astrocytes provide significant neuroprotection and thus making astrocytic SRF a potential therapeutic target. In addition, the identification of SRF-dependent genes and pathways and the elucidation of neuroprotective mechanisms may open the door for novel pharmacologic therapies aimed at converting astrocytes to a neuroprotective state and allowing for optimized neuronal recovery following injury or in the setting of neurodegenerative disease.

## Materials and Methods

### Animals

*Srf*-floxed mice were previously described (Ramanan et al., 2005). These mice were bred with GFAP-Cre (generously provided by Dr. David Guttmann, Washington University School of Medicine, St. Louis, MO) and GFAP-Cre-ERT (generously provided by Dr. Suzanne J. Baker, St. Jude Children’s Research Hospital, Memphis, TN) transgenic mice to obtain *Srf*^f/f; GFAP-Cre+/-^ (*Srf*^GFAP^CKO) and *Srf*^f/f;GFAP-CreERT+/-^ (*Srf*^GFAP-ER^CKO) respectively. *Srf*^f/f^ mice served as control in all experiments. Control and mutant mice were housed together, and cage mates were randomly assigned to experimental groups. All experiments were conducted blinded to the genotype of the mice used. All the procedures in this study were performed according to the rules and guidelines of the Committee for the Purpose of Control and Supervision of Experimental Animals (CPCSEA), India. The research protocol was approved by the Institutional Animal Ethics Committee (IAEC) of the Indian Institute of Science.

### Tamoxifen treatment

Tamoxifen (Alfa Aesar, Cat. No. J63509-03) was dissolved in corn oil (MP Biomedicals, Cat. No. 0290141405) at 30 mg/ml. This was injected intraperitoneally to 6-8-week-old *Srf*^GFAP-CreER^CKO mice at 9 mg/40g of body weight for five consecutive days.

### Immunohistochemistry

Mice were fixed by transcardial perfusion using 4% PFA. The brains were cryoprotected in 30% sucrose, frozen and stored in −80° C until further use. For staining, 30 µm thick cryosections were incubated in the blocking/permeabilization solution containing 0.3% Triton-X and 3% goat serum in 1X PBS (pH 7.4) for 1 hour followed by overnight incubation in the primary antibody. The brain sections were then washed in PBS and incubated in the secondary antibody for 1 hour. The sections were finally mounted in a DAPI-containing mounting medium (Vector Laboratories). The following primary antibodies were used: mouse anti-GFAP (1:1000; Sigma, #G-145), rabbit anti-GFAP (1:1000; Z0334, Dako), goat anti-Sox9 (1:500, R&D Systems, #AF3075), mouse anti-Vimentin (1:50; DSHB, #40E-C), rabbit anti-S100β (1:1000; Sigma, #S2644), rabbit anti-Iba1 (1:1000; Wako, #019-19741), mouse anti-NeuN (1:1000; Chemicon, #MAB-377), chicken anti-β-gal (1:1000; Aves, #BGL-1040), rabbit anti-PhosphoHistoneH3 (1:500; Sigma, #H0412), guinea pig anti-Piccolo (1:400, Synaptic systems, #142104), mouse anti-GluA1 (1:200, Synaptic systems, # 182011), rabbit anti-Tyrosine hydroxylase (1:500, Merck, #AB152), and anti-MOAB2 (1:500, Novus Bio, #NBP2-13075). The following secondary antibodies were used: AlexaFluor-488 and −594-conjugated anti-rabbit, anti-chicken, anti-guinea pig and anti-mouse at 1:1000 dilution (Life Technologies). Biotinylated anti-mouse and anti-rabbit secondary antibodies (1:250; Vector Laboratories) were used along with Vectastain ABC kit and ImmPACT VIP kit (Vector Labs). Images were captured using fluorescence microscope (Eclipse 80i, Nikon), or confocal microscope (LSM 880, Zeiss).

### FluoroJade-C staining

FluoroJade-C (FJC) (#1FJC, Histo-Chem Inc.) staining was carried out according to the manufacturer’s instructions. Briefly, 35 µm thick cryosections were mounted on Superfrost plus slides (Brain Research Laboratories) and allowed to dry at 60 °C for 60 min. Slides were immersed in 80% EtOH with 1% NaOH for 5 min, followed by 2 min in 70% EtOH, 2 min in distilled water, and incubated in 0.06% potassium permanganate solution for 10 min. Slides were subsequently rinsed in water, transferred to a 0.0001% solution of FJC in 0.1% acetic acid. The slides were finally rinsed in distilled water, air dried, cleared in xylene and mounted with DPX mountant. Fluorescence imaging was carried out using an epifluorescence microscope (Eclipse 80i, Nikon) with appropriate excitation/emission filters and captured using Metamorph Software.

### Black-Gold II staining

Black-Gold II staining was done using the Black-Gold II myelin staining kit (AG105, Merck Inc, USA) as per the manufacturer’s protocol. Briefly, Black-Gold II powder was resuspended in 0.9% NaCl (Saline solution) to a final concentration of 0.3% and sodium thiosulphate solution was diluted to 1% with MilliQ water freshly before use. 0.3% Black-Gold II and 1% sodium thiosulphate were prewarmed to 60 °C. 35 µm floating brain sections were rehydrated in 1X PBS for 2 minutes and then transferred to a pre-warmed Black-Gold II solution. The sections were incubated in Black-Gold II solution at 60 °C for 20 minutes and then transferred to 1X PBS and rinsed twice for 2 minutes each at room temperature. The sections were then incubated with 1% sodium thiosulphate solution at 60 °C for 3 minutes to stop the reaction. The sections were then rinsed with 1X PBS thrice for two minutes each at room temperature. The sections were then mounted on positively charged slides and allowed to dry overnight at room temperature. Next day, the slides were dehydrated using a series of graded alcohols, cleared in xylene for 1 minute and coverslipped with DPX mounting media. All the Black-Gold II brightfield images were taken on a Nikon Eclipse 80i upright microscope and analyzed using the ImageJ software. For Black-Gold II intensity measurements, the images were converted into 8-bit, binarized and inverted using the invert LUT function in ImageJ. Following this, intensity measurements were made with ROIs of suitable sizes (200 x 200 µm^2^).

### Quantification of fluorescence intensity and cell numbers

For measuring fluorescence intensity, images were scaled for 10X magnification. Ten to twelve areas of field in the same rostro-caudal axis were chosen per image, and the intensities were measured using ImageJ after subtracting the background fluorescence from both control and knockout sections. For the hippocampus, cell numbers or fluorescence intensities were measured in the *stratum oriens* and *stratum radiatum*. For cell counts, images were taken at 10X magnification. Four ROIs in the same rostro-caudal axis were chosen per image, and the number of cells per ROI were counted with the cell counter plugin using ImageJ and area was converted to mm^2^.

### Sholl analysis

To measure the hypertrophy of reactive astrocytes, semi-automated Sholl analysis plugged in ImageJ was used. First, the astrocytes and processes of interest were outlined to exclude adjacent cells or areas. Templates of concentric circles from 10-100 µm (increasing by 10µm) were overlaid from the center of the cell soma. For each cell, densitometric thresholds were set to subtract the background, followed by converting to mask, Despeckle and skeletonize the image to analyze the particles. Measurements obtained from each individual cell were recorded. Two-way ANOVA with Sidak’s *post hoc* test and mean difference was calculated.

### BBB Permeability Assay

Assay to measure the integrity of the BBB was performed as previously described (Andreone et al., 2017). Briefly, 17-mon old *Srf*^GFAP-ER^CKO (15-mon post-Tam injection) mice were deeply anesthetized with isoflurane and injected with 20 µl of 10 kDa dextran fluorescein (Invitrogen; #D1820) into the left cardiac ventricle, and allowed to circulate for 5 min. Their brains were collected and post-fixed in 4% PFA overnight, frozen and stored at −80 °C. 30 µm thick cryosections were mounted using mounting media supplemented with DAPI (Vector Labs) and analyzed using an epifluorescence microscope (Eclipse 80i, Nikon). Brain sections from the similar rostro caudal (Bregma) position were analyzed. At least 12 different regions were taken and the ratio of the fluorescence intensity (inside versus outside the vessel) was measured.

### Standard qRT-PCR

Quantitative RT-PCR was done with 100 ng of cDNA and KAPA SYBR FAST ABI prism kit (Cat. No. KK4604) using the following program: 95 °C or 3 min followed by 39 cycles of 95 °C for 5 sec, 55 °C for 30 sec and 72 °C for 40 sec. The PCR reaction was carried out in QuantStudio 7 Flex Real-Time PCR System (Invitrogen Biosciences). All the primer sequences were published previously (Liddelow et al., 2017).

### Quantification of the number of synapses

Neocortical sections (30 µm) from tamoxifen injected *Srf*^GFAP-ER^CKO mice and control littermates were immunostained for Piccolo (presynaptic marker) and GluA1 (postsynaptic marker). The images were captured using a confocal microscope (Zeiss LSM880 AxioObserver; PlanApo 63X/1.4 oil objective) with a lateral sampling of 40 nm and an axial sampling of 1 µm. The area of imaging was optimized as 1024 x 1024 pixels and the illumination and imaging conditions (10% power of the lasers, sampling, and pinhole size of 1 Airy unit) were kept constant between the samples. The samples were exported to MetaMorph image analysis module (Molecular Devices, USA) for further segmentation and analysis. The imported multi-wavelength image stacks in 3D were separated into individual channels and a maximum intensity projection was made for each wavelength channel labeling for pre- and postsynaptic markers. The average intensity and standard deviation of each image was obtained through an inbuilt module to quantify image statistics in MetaMorph. The threshold of the images was set at an intensity of (average + standard deviation) for each image. The regions which were between 0.2 µm and 2 µm were automatically detected and filtered to create a binary mask for each image. Clusters with total area less than that of 25 pixel^2^ (1 pixel = 40 nm) with 0.2 µm (length and breadth) were discarded. The filtering parameters of the thresholded image were chosen to match with the existing information on the size of pre- and postsynaptic compartments obtained through electron microscopy (Ippolito and Eroglu, 2010). The binarized images of presynaptic and postsynaptic markers were then analyzed for an overlap for 1 or more pixels between them using the integrated morphometry analysis module in MetaMorph. An overlap of one or more pixels was identified to indicate the presence of a structural synapse. The number of overlapping puncta was counted for each overlaid binary image of pre- and postsynaptic markers. The area of the field for counting synapses was 100×100 µm^2^.

### Quantification of β-amyloid plaques

The stained brightfield sections were imaged in a Nikon 80i upright microscope. The data were collected as 16-bit images and saved as TIFF format using Metamorph (Molecular Devices). The 16-bit color images were converted into inverted 16-bit monochrome TIFF files and subsequentially used for automated estimation of β-amyloid plaques. The data obtained were pseudo-labelled and blind analyses were performed by a person different from the one who imaged the samples. For every brain section, cortex and hippocampus were marked with randomly chosen regions of interest (ROIs). The ROIs were marked such that in each hemisphere cortex had 2 ROIs of 500 x 500 pixel^2^ and hippocampus had three randomly chosen regions 200 x 200 pixel^2^ in the *stratum radiatum*. The images were then auto thresholded for dark objects. The Average and Gray level value of thresholded regions were estimated and half this Average Gray level value obtained after initial thresholding was taken as the cut-off intensity for detection of β-amyloid plaques. The thresholded objects were measured using integrated morphometry analysis and objects larger than a pixel area of 100 pixel^2^ were automatically chosen and counted in each region of interest. The number of plaques in the cortical and hippocampal regions in one brain section were averaged to obtain a mean distribution of plaques across all regions per sample.

### Preparation of hippocampal slices

Acute brain slices were prepared from tamoxifen-treated male and female *Srf*^f/f^ (control) and *Srf*^GFAP-ER^CKO mice at 3-mon and 15-mon post-tam administration. Mice were sacrificed by cervical dislocation, and brains were isolated and kept in ice-cold sucrose-based artificial cerebrospinal fluid (aCSF; cutting solution) comprising (mM): 189 Sucrose (Sigma #S9378), 10 D-Glucose (Sigma #G8270), 26 NaHCO_3_ (Sigma #5761), 3 KCl (Sigma #P5405), 10 MgSO_4_.7H_2_O (Sigma #M2773), 1.25 NaH_2_PO_4_ (Sigma #8282) and 0.1 CaCl_2_ (Sigma #21115). After 30 seconds, the brain was glued to the brain holder of the vibratome (Leica #VT1200), 350 µm thick horizontal slices were prepared, and the cortex was dissected to isolate the hippocampus. All the slices were kept in slice chamber containing aCSF comprising (mM): 124 NaCl (Sigma #6191), 3 KCl (Sigma #P5405), 1 MgSO_4_.7H_2_O (Sigma #M2773), 1.25 NaH_2_PO_4_ (Sigma #8282), 10 D-Glucose (Sigma #G8270), 24 NaHCO_3_ (Sigma #5761), and 2 CaCl_2_ (Sigma #21115) in water bath (Thermo Scientific #2842) at 37°C for 40-45 minutes. Following recovery, slices were maintained at room temperature (RT) until transferred to a submerged chamber of ∼1.5 ml volume in which the slices were perfused continuously (2-3 ml/minute) with warmed (34°C) and carbogenated aCSF. Post dissection, every step was carried out in the presence of constant bubbling with carbogen (2-5% CO_2_ and 95% O_2_; Chemix). All measurements were performed by an experimenter blind to the experimental conditions.

### Extracellular Field Recordings

Field excitatory postsynaptic potentials (fEPSP) were elicited from pyramidal cells of the CA1 region of *stratum radiatum* by placing concentric bipolar stimulating electrode (CBARC75, FHC, USA) connected to a constant current isolator stimulator unit (Digitimer, UK) at CA3 of Schaffer-collateral commissural pathway and recorded from *stratum radiatum* of CA1 area of the hippocampus, with 3-5 MΩ resistance glass pipette (ID: 0.69mm, OD: 1.2mm, Harvard Apparatus) filled with aCSF. Signals were amplified using an Axon Multiclamp 700B amplifier (Molecular Devices), digitized using an Axon Digidata 1440A (Molecular Devices), and stored on a computer using pClamp10.7 software (Molecular Devices). Stimulation frequency was set at 0.05 Hz. I/O were obtained by setting a stimulation range of 20-40 µs and by adjusting the stimulus intensity by 10 µA per sweep with increments from 0-300 µA. Paired pulse ratio (PPR), which is a cellular correlate of release probability of neurotransmitters, was assessed with a succession of paired pulses separated by intervals of quarter log units, with the interval ranging from 3 to 1000 ms and were delivered every 20s. The degree of facilitation (CA3-CA1) was determined by taking the ratio of the initial slope of the second fEPSP relative to the first fEPSP. PPR >1 was considered as a paired pulse facilitation.

Long term potentiation (LTP) was induced using a theta burst protocol (TBS). A baseline period of 15-minute basal fEPSP was recorded at a stimulation intensity that elicited an approximately half-maximal response. Stimulation intensity remained constant throughout the experiment, including during TBS and 60 minutes post potentiation. Following the baseline period, the TBS protocol was delivered consisting of five bursts (10 stimuli at 100 Hz) at 5 Hz (theta frequency), repeated four times at an interval of 20 s (Booth et al., 2014). fEPSP after TBS were recorded for 60-minute. Slices with high FV/fEPSP ratio, and unstable for 15-minute baseline were discarded from the analysis.

Data analyses were performed using Clampfit 10.7 and Excel 2016. Data are represented as Mean ± SEM. Student unpaired *t-*test were performed to determine the statistical significance. To determine statistical difference in I/O, linear regression was performed on the individual slices to determine the slope of the relationship, and then one-way *ANOVA* (genotype) was performed on the regression points (Zaman et al., 2000). Example traces are those recorded for 1-2 minute around the time point indicated in the graph.

### Kainic acid (KA) injection

Intracerebroventricular (ICV) injection of KA into the brain was performed as described earlier (Jin et al., 2009). Briefly, male, and female *Srf*^GFAP-ER^CKO (4 months post-Tam injection) and control littermates were deeply anaesthetized with 3% isoflurane (Sosrane, NEON, Mumbai, India) for 15 min. The anesthetized animal was placed on a stereotaxic apparatus (KOPF, TUNJUNG, CA, USA) with continuous administration of 1.5 - 2% isoflurane for 1h with vehicle air. Mice were injected with 4 µl saline containing 0.1 – 0.2 µg of KA using a 10 µl Hamilton syringe fitted with a 28-gauge needle (Hamilton Company, Nevada, USA). The needle was inserted through a hole perforated on the skull (using a dental drill), into the right lateral ventricle using the following coordinates (in mm with reference to bregma): anteroposterior (AP), −0.2; mediolateral (ML), −2.9; dorsoventral (DV), −3.7. After 5 min, the needle was withdrawn over 3 min to prevent backflow. The mice were warmed under infrared (IR) lamp (245 V, 150 W) placed 1-2 feet away until being awakened. After KA injection, animals were carefully monitored for 2h and seizure activity was scored by the following Racine score: (0) normal activity, (+1) rigid posture/ immobility, (+2) head bobbing, (+3) forelimb clonus and rearing, (+4) rearing and falling, (+5) tonic-clonic seizures, and (+6) death within 2h. The animals were monitored for 7 days and then transcardially perfused. The brains were isolated, post-fixed overnight in 4% PFA, cryoprotected in 30% sucrose, frozen and sectioned at 30 µm in a cryostat (Leica, CM 1850).

### TUNEL assay

The TUNEL assay was carried out using Click-IT Plus TUNEL assay kit (Molecular Probes, Thermo Fisher Scientific). Briefly, 30 µm paraformaldehyde-fixed cryosections were permeabilized with proteinase K solution for 15 min and then incubated with TdT reaction mixture for 60 min at 37 °C and subsequently with FITC-labeled dUTP for 30 min. The slides were washed with 3% BSA in PBS for 5 min and rinsed in 1X PBS. The slides were mounted using a mounting medium containing DAPI (Vector Labs) and observed using an epifluorescence microscope (Eclipse 80i, Nikon) using appropriate filters and captured using Metamorph software. Numbers of TUNEL positive cells in the CA1, CA3 and DG regions of entire rostral to caudal brain regions were counted using Image J software. The area of the field for counting the number of TUNEL+ cells was 250×250 µm^2^ and converted to mm^2^.

### 6-Hydroxydopamine (6-OHDA) injections

6-OHDA (HelloBio, UK, #HB1889) injections in mice were performed as described earlier (Grealish et al., 2010). Briefly, the mice, *Srf*^GFAP-ER^CKO (9 mpi) and control littermates were anesthetized with continuous administration of 1.5 - 2% isofluorane (Sosrane, NEON, Mumbai, India) and placed on the stereotaxic apparatus (KOPF, TUNJUNG, CA, USA). 6-OHDA injections were made in the substantia nigra pars compacta (SNpc) using 10 µl Hamilton syringe fitted with a 28-gauge needle (Hamilton Company, Nevada, USA). The 6-OHDA toxin used was at a concentration of 1.6 µg/µl prepared in ascorbic acid solution (0.2mg/ml of ascorbic acid in 0.9% saline, filter sterilized). From this, each mouse received a total volume of 1.5 µl using the following stereotaxic coordinates: anteroposterior (AP) = 3.0, Mediolateral (ML) = 1.2, and dorsoventral (DV) = 4.5, with a flat skull position (coordinates in mm with reference to Bregma). Injections were made at 0.5 µl/min with an additional 5 min to allow the toxin to diffuse and 3 min for slow withdrawal of the syringe. Following injections, the mice were warmed under an infrared (IR) lamp (245 V, 150 W) placed 1-2 feet away until being awakened and returned to the home cage. The animals were monitored for 10 days and then transcardially perfused. The brains were isolated, post-fixed overnight in 4% PFA, cryoprotected in 30% sucrose, frozen and 30 µm sections were prepared in a cryostat.

### Open Field Test

Open field test was used to assess baseline locomotion. The open field test was done under ambient lighting in a 45 (L) x 45 (W) x 35 (H) cm square acrylic box. Mice were gently introduced in the center of the arena and were allowed to explore freely for 10 minutes. The behavior was recorded using an overhead webcam (Logitech C270HD) at 720p and 15 fps. Between each test, the arena was thoroughly cleaned with 70% ethanol to remove any odor cues. Movement of mice was tracked using custom MATLAB® code developed in our lab (www.github.com/swanandlab). Total distance traveled (cm) was automatically measured using the obtained coordinates.

### Fear conditioning

Declarative memory was tested using the contextual fear conditioning paradigm. The mice were handled for 4 days prior to training to reduce any anxiety-related effect on their memory performance. Two days prior to training, mice were habituated to transportation and room environments for 10 minutes. Configuration of sound attenuated contexts were as follows: context A – floor made up of steel grids, white light, 15% acetone; context B – triangle shed, laminated white sheet as floor to provide different tactile cues,75% ethanol.

#### Training

On training days, fear conditioning chambers were cleaned with 15% acetone. The tray below was cleaned and sprayed before placing any animal in the chamber. Animals were allowed to explore context A for 90 seconds and then a 2.8kHz 75db tone was played for 30 seconds. Animals received a 0.7mA mild foot-shock in the last 2 seconds of 30 second 2.8kHz 75 dB tone in context A. This was repeated twice with a time interval of 15 seconds. After the last shock, animals were taken out of the chamber and brought back to their home cage.

#### Recent memory recall test in context A

24h post-training, animals were scored for their freezing response as a parameter of their memory. The first 90 seconds of freezing responses were scored manually with a custom scoring assistant plugin in ImageJ. A minimum of 1.2 second continuous freezing bouts were taken as a freezing response. Manually recorded data was analyzed using MATLAB.

#### Recent memory recall test in context B

48h post-training, animals were scored in context B for their freezing responses for the first 90 seconds.

#### Remote memory recall

One month after the fear conditioning animals were tested again in context B, as explained above, and scored for their freezing response. 24 hours later, the animals were tested in context A and freezing responses were recorded. In the remote recall, the order of contexts was inverted to avoid the CS-US dissociation.

### Barnes maze

Barnes maze test was performed as previously described (Martyn et al., 2012). A white circular platform, 92 cm in diameter, with 20 equally spaced 5 cm-wide holes along the periphery was used. The platform was elevated 95 cm from the floor. The behavior was recorded using an overhead webcam (Logitech C270HD) at 720p and 15 fps. A dark escape chamber was placed under one of the holes, called the target hole, which was indistinguishable from other holes. Three different visual cues, triangular, square, and circular in shape, were placed 10 cm from the edge of the platform, 120° apart from each other for mice to orient themselves in space. Cues remained constant throughout training and testing. Uniform illumination of the platform by bright LED light created an aversive stimulus to motivate the mice to seek the target hole. Before every trial, the arena was cleaned with 70% ethanol to eliminate olfactory cues. In case of unsuccessful trials, mice were gently guided to the target hole. One day before the training trial, the mice were habituated to the maze by placing them in a clear transparent plastic cylinder (15 cm in diameter) in the middle of the platform for 30 seconds, following which they were guided to the target hole. Mice were given 3 minutes to enter the escape chamber. The mice that did not enter the escape chamber were gently nudged with the cylinder to enter the chamber. Mice spent 1 min in the escape chamber before being returned to their home cage. Habituation consisted of 2 trials spaced 4 hours apart. On training day, mice were placed in an opaque plastic cylinder covered by an opaque lid (15 cm in diameter) for 15 seconds to randomize starting orientation. Each trial was initiated by lifting the cylinder and the trial lasted 2 minutes. The trial was stopped if the mouse entered the escape chamber. If the mouse did not get into the escape chamber within 2 minutes of trial, it was gently guided to the target hole with the clear plastic cylinder. Mice were returned to their home cage after 15 secs of entering the escape chamber. Training trials were performed twice daily, with 1 hour interval between them with a pseudo-random starting location in the four quadrants. Trials were conducted for 4 consecutive days, until the learning curve leveled off. The test was conducted 2 hours after the last training trial. The escape chamber was removed for the probe test and the mice were allowed to explore the maze for 1 minute. The videos were later analyzed for trajectories and other parameters using custom MATLAB scripts (www.github.com/swanandlab). Time to reach the correct location of the target hole was reported as latency to the target hole when the body centroid came within 10 cm of the target hole. Time spent in the target quadrant was also measured and reported.

### Astrocyte isolation for transcriptome analysis

Astrocytes were isolated by magnetic assisted cell sorting (MACS) approach using the anti-ACSA2 microbeads (Miltenyi Biotec Inc., Germany). Briefly, the brains were isolated from 5–6-week-old *Srf*^GFAP^CKO mice and meninges were removed in ice-cold homogenization buffer (For 10 ml: 1 ml 10X HBSS w/o Ca^2+^ and Mg^2+^, 1 µl of 1mg/ml DNase I, 150 µl of 1M HEPES, 120 µl of D-glucose, 9 ml milli-Q water). The neocortex and hippocampus were isolated and homogenized in a glass potter with pestle (7 ml volume) in 5 ml of ice-cold homogenization buffer. The homogenate was strained through a 70 µm cell strainer and centrifuged at 300 x *g* for 10 min at 4 °C. The cell pellet was resuspended in 37% Percoll in 15-ml centrifugation tube and centrifuged at 800 x *g* for 20 mins at 4 °C. To make 37% Percoll, first a 100% Percoll solution was prepared by mixing 9 parts of Percoll with 1 part of 10X DEPC-treated PBS (DPBS). This 100% Percoll was diluted to 37% using 1X DPBS. The myelin in the top layer was carefully removed and discarded. The remaining supernatant was transferred to a fresh 50-ml centrifugation tube and 3 volumes ice-cold PB buffer (0.5% BSA in 1X DPBS) to dilute the Percoll. The cell pellet was gently resuspended in 5 ml ice-cold PB buffer. Both these tubes were centrifuged at 700 x *g* for 10 min at 4 °C. The supernatant was discarded, and the cell pellets were pooled together in 80 µl PB buffer and 10 µl of FcR blocking reagent was added per 10^7^ cells. After 10 min incubation at 4 °C, 10 µl of anti-ACSA2 microbeads were added per 10^7^ cells and incubated for 15 min at 4 °C. The ACSA2+ astrocytes were isolated using the midi-MACS separator and LS columns (Miltenyi Biotec) as per the manufacturer’s protocol. The ACSA2+ astrocytes were eluted from the magnetic columns and pelleted by centrifugation at 300 x *g* for 10 min at 4 °C and resuspended in RNA Later (Invitrogen).

### RNA sequencing and analysis

The sequence data was generated using Illumina HiSeq 2500. Data quality was checked using FastQC and MultiQC software. The data was checked for base call quality distribution, % bases above Q20, Q30, %GC, and sequencing adapter contamination. All the samples passed QC threshold of Q_20_>95%. Raw sequence reads were processed to remove adapter sequences and low-quality bases using fastp. The QC passed reads were mapped onto indexed Mouse reference genome (GRCm 38.90) using STAR v2 aligner. On average 99.03% of the reads aligned onto the reference genome. Gene level expression values were obtained as read counts using featureCounts software. Expression similarity between biological replicates was checked by spearman correlation and Principal Components Analysis. For differential expression analysis the biological replicates were grouped as Control and Test. Differential expression analysis was carried out using the edgeR package after normalizing the data based on trimmed mean of M (TMM) values. After normalization 27540 features (52.32%) have been removed from the analysis because they did not have at least 1 counts-per-million in at least 4 samples. Genes with absolute log_2_ fold change ≥ 1.5 and adjusted p-value ≤ 0.05 were considered significant. The expression profile of differentially expressed genes across the samples is presented in volcano plots and heatmaps. The genes that showed significant differential expression were used for Gene Ontology (GO) and pathway enrichment analysis using DAVID (https://david.ncifcrf.gov/).

### Statistics

Analyses were done using GraphPad Prism. The comparisons between two groups were done using unpaired two-tailed Student’s *t*-test or one-way ANOVA with Tukey *post hoc* test. For 4 group comparisons with 2 variables (context and genotype) in the fear conditioning test, 2-way ANOVA was used followed by Sidak’s *post hoc* test to analyze the interactions between the 2 variables. The standard error mean (SEM) was calculated and depicted as error bars in the graphs. All the statistical details for each experiment, including the *n* value, the statistical test used, P value, significance of comparisons are mentioned in the figure legends.

## Acknowledgments

We thank Dr. Suzanne Baker (St. Jude’s Hospital, Memphis, TN, USA) for generously sharing the GFAP-ERT transgenic mouse line; Dr. Joseph LoTurco (Univ. of Connecticut, Storrs, CT, USA) for sharing the piggyBac plasmids. Dr. Hiyaa Ghosh (NCBS, Bangalore) and Dr. Bhavana Muralidharan (InStem, Bangalore) for sharing antibodies; We thank the Bioimaging Facility and the Central Animal Facility for confocal imaging and animal care, respectively.

## Author contributions

Conceptualization: NR; Methodology: BJ, DN, JPC, SM and NR; Investigation: SCRT, MJ, SS, SD, VV, AN, DN and NR; Visualization: SCRT, MJ, SS, SD, VV, AN and NR; Supervision: NR; Writing—original draft: NR; Writing—review & editing: SCRT, MJ, SS, SD, VV, AN, DHG, BJ, DN, JPC, SM and NR.

## Data and materials availability

All data are available in the main text or the supplementary materials.

