## Supplemental Material for "SRF-deficient astrocytes provide neuroprotection in mouse models of excitotoxicity and neurodegeneration"

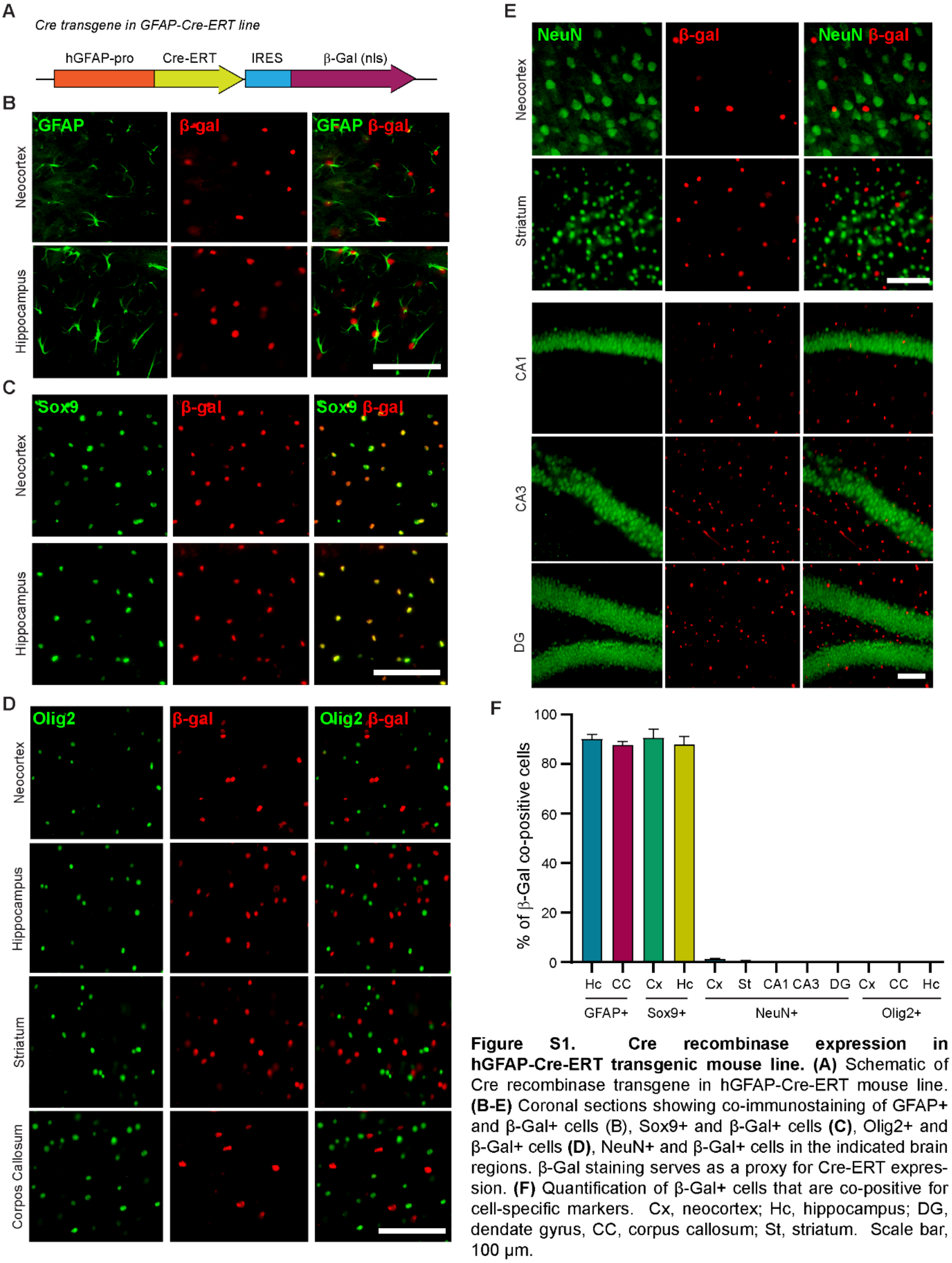

**Figure S1. Cre recombinase expression in hGFAP-Cre-ERT transgenic mouse line. (A)** Schematic of Cre recombinase transgene in hGFAP-Cre-ERT mouse line. **(B-E)** Coronal sections showing co-immunostaining of GFAP+ and β-Gal+ cells (B), Sox9+ and β-Gal+ cells **(C)**, Olig2+ and β-Gal+ cells **(D)**, NeuN+ and β-Gal+ cells in the indicated brain regions. β-Gal staining serves as a proxy for Cre-ERT expression. **(F)** Quantification of β-Gal+ cells that are co-positive for cell-specific markers. Cx, neocortex; Hc, hippocampus; DG, dendate gyrus, CC, corpus callosum; St, striatum. Scale bar, 100 µm.

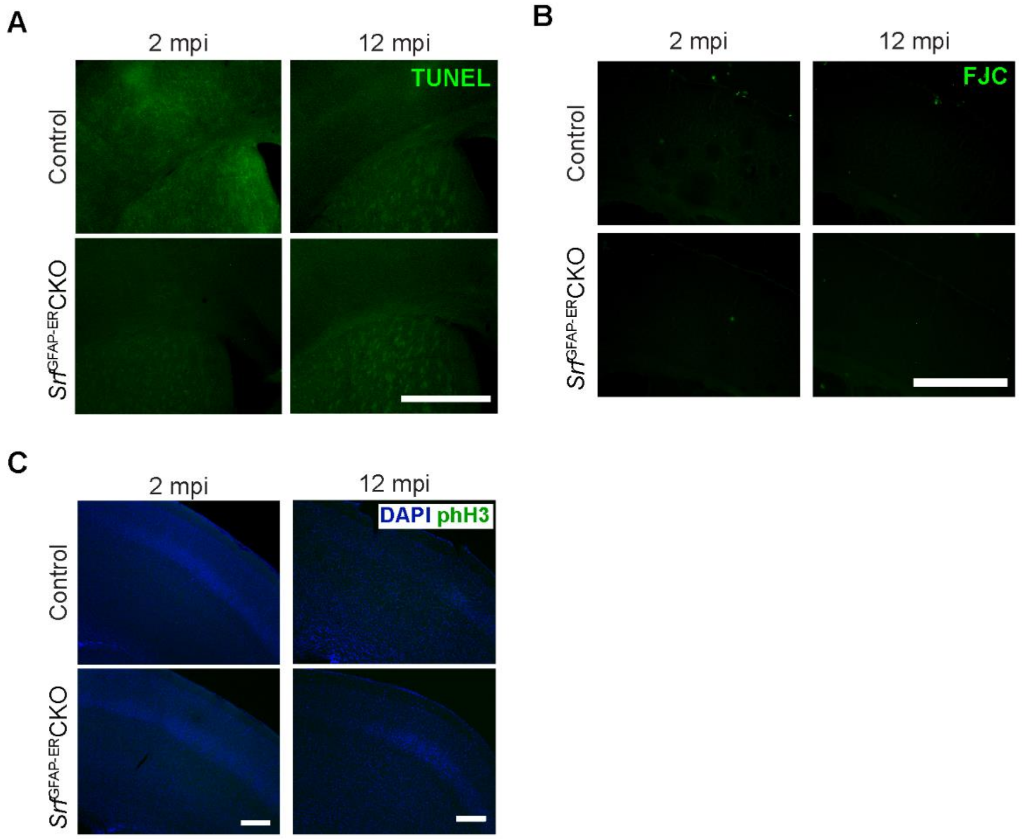

**Figure S2. Absence of proliferation and cell death in *Srf*$^{GFAP\text{-}ER}$CKO mice. (A)** Coronal section showing TUNEL staining in the neocortex of control and *Srf*$^{GFAP\text{-}ER}$CKO mice at 2 mpi and 12 mpi. **(B)** Coronal sections showing FluoroJade-C (FJC) staining of the neocortex. **(C)** Coronal sections showing immunostaining for phosphor-Histone H3 (phH3) in the neocortex. DAPI labels the nuclei. Scale bar, 100 µm.

Figure S3

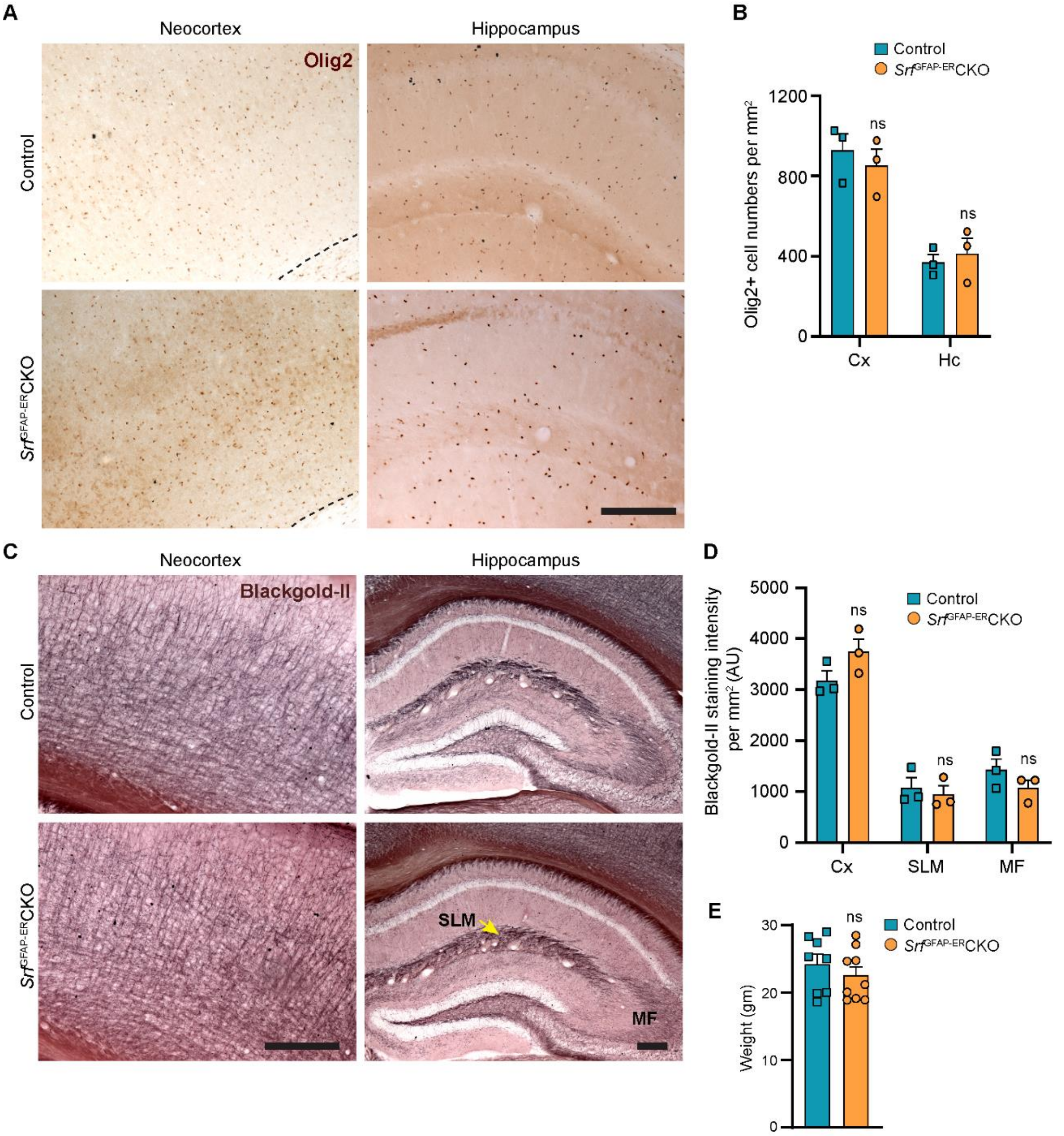

**Figure S3. Oligodendrocyte lineage cells and myelination in *Srf*$^{GFAP-ER}$CKO mice. (A)** Coronal sections showing immunostaining for Olig2 in neocortex and hippocampus in control and *Srf* mutant mice at 12 mpi. Scale, 200 μm. **(B)** Quantification of number of Olig2+ cells in (A). n=3 mice. **(C)** Black-Gold II myelin staining of coronal sections of control and *Srf* mutant mice at 12 mpi. Scale, 200 μm. **(D)** Quantification of myelin staining intensity from the neocortex (Cx), *Stratum lacunosum moleculare* (SLM) and mossy fibers (MF) and in (C). n=3 mice. **(E)** Measurement of body weights of control and SRF mutant mice at 12 mpi. No difference in body weights was seen between the two groups. n=8-9 mice. ns, not significant. Unpaired t-test. Data are represented as mean ± SEM. AU, arbitrary units.

Figure S4

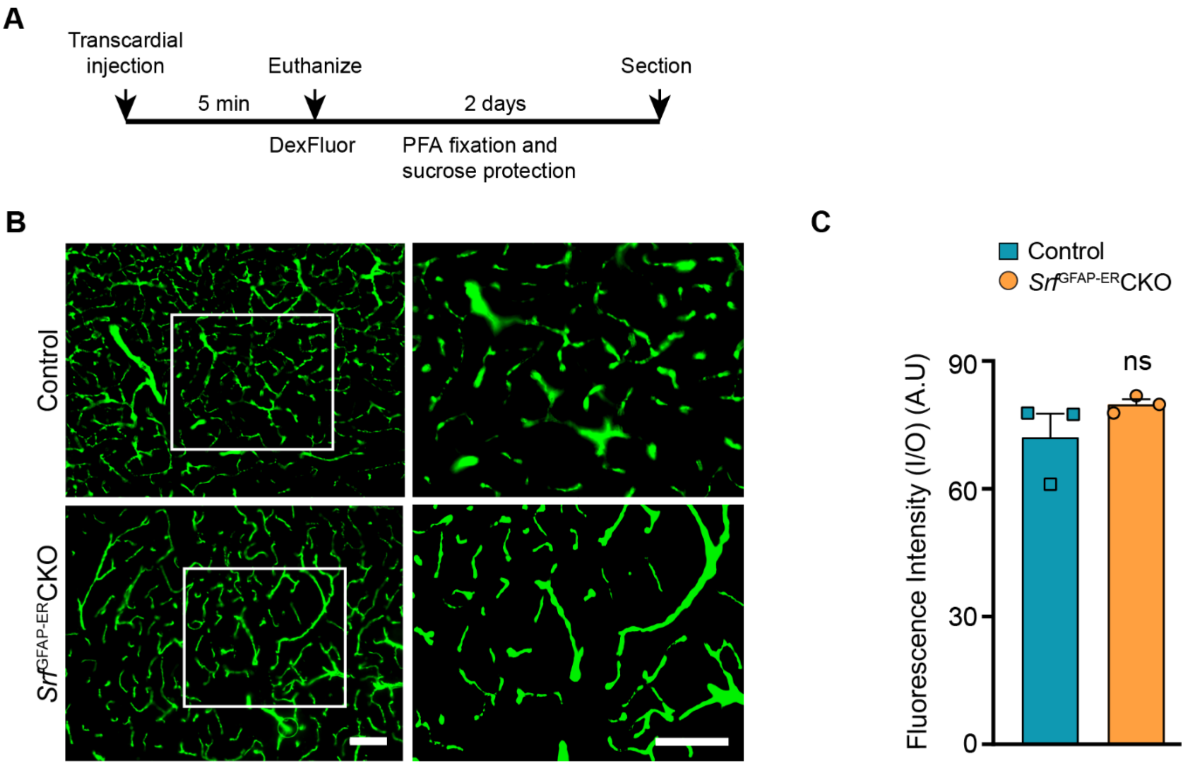

**Figure S4. BBB integrity is not compromised in *Srf*$^{GFAP\text{-}ER}$CKO *mice* mutant mice. (A)** Schematic of dextran fluorescein (DexFluor) injection. **(B)** Visualization of dextran fluorescein (10 kDa) in coronal sections of the neocortex. **(C)** Quantification of ratio of fluorescence intensity inside versus outside of blood vessels in (B). n=3 mice. ns, not significant. Unpaired t-test. Data are represented as mean ± SEM. Scale bar, 50 μm. AU, arbitrary units.

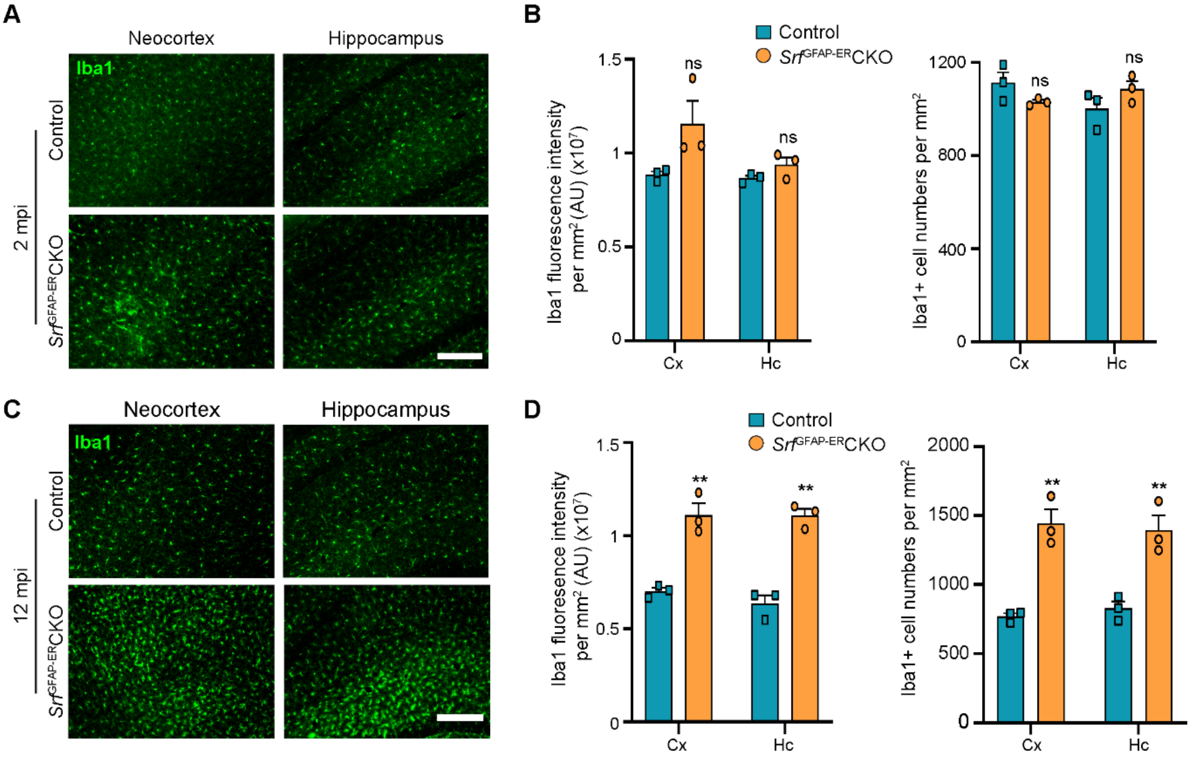

**Figure S5. Increase in Ibal intensity and cell numbers in older but not younger *Srf* mutant mice. (A)** Immunostaining of coronal sections showing Iba1+ microglia in the neocortex and hippocampus in control and mutant mice at 2 mpi. **(B)** Quantification of Iba1+ cell number and relative Iba1 fluorescence intensity in the neocortex and hippocampus shown in A. **(C)** Immunostaining of coronal sections showing Iba1+ microglia in the neocortex and hippocampus in control and mutant mice at 12 mpi. **(D)** Quantification of Iba1+ cell number and relative Iba1 fluorescence intensity in the neocortex and hippocampus shown in B. mpi, months post-tamoxifen injection, Cx, neocortex; Hc, hippocampus; St, striatum. AU, arbitrary units. n=3 mice. Data are represented as mean ± SEM. * $P < 0.05$, ** $P < 0.01$, ns, not significant. Unpaired t-test. Scale bar, 100 µm.

Figure S6

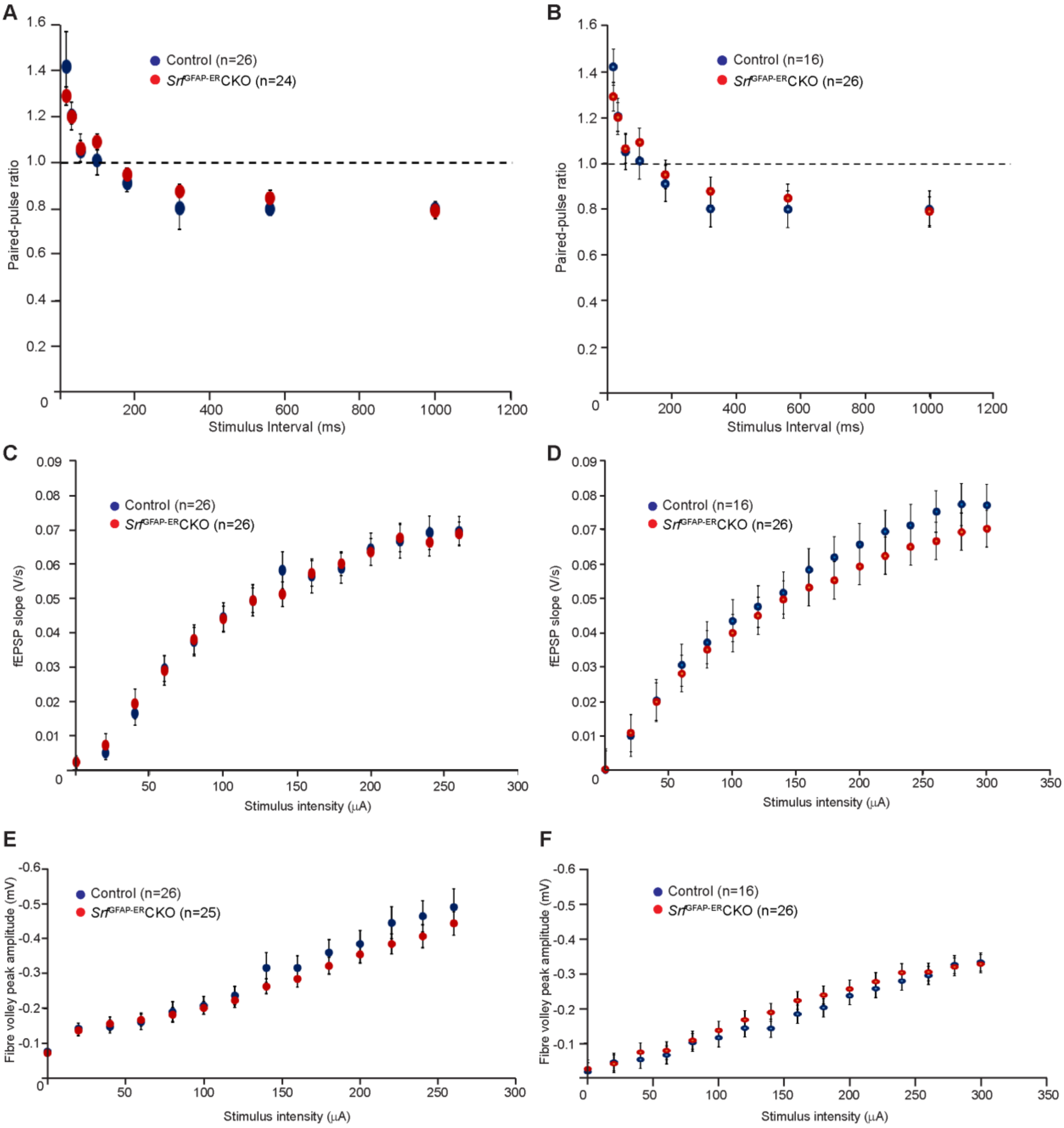

**Figure S6. Normal synaptic functions in *Srf*$^{GFAP\text{-}ER}$CKO mice.** Basal synaptic properties in the Schaeffer-Collateral pathway in hippocampus were measured in *Srf*$^{GFAP\text{-}ER}$CKO mice and control littermates at 3 mpi **(A, C, E)** and 15 mpi **(B, D, F)**. Analyses of paired-pulse ratio (A, B), post-synaptic response to stimulus intensity (C, D) and summated action potential (E, F) did not show any significant difference between the two groups of mice. The number of slices recorded from are indicated in parentheses. (n=8 control; n=5 mutant mice per group). Unpaired t-test. Data are represented as mean ± SEM.

Figure S7

**A**

| Gene name | ENTREZ ID | Padj. value | Log2FC |
|---|---|---|---|
| ***Immune response and innate immunity*** | | | |
| Cd68 | 12514 | 3.71519E-08 | 4.307 |
| H2-T23 | 15040 | 1.15514E-05 | 3.551 |
| Cd84 | 12523 | 2.91635E-19 | 5.351 |
| Lst1 | 16988 | 3.72083E-10 | 4.009 |
| Ifit3 | 15959 | 1.61217E-06 | 3.142 |
| Trim30a | 20128 | 1.66639E-05 | 2.996 |
| Trim30b | 244183 | 7.25273E-11 | 4.07 |
| ***Inflamamatory response*** | | | |
| *Nlrp1b* | 637515 | 4.59748E-11 | 3.378 |
| *Ccl12* | 20293 | 3.74557E-16 | 7.759 |
| *Cxcl13* | 55985 | 1.26903E-12 | 6.821 |
| *Ccl5* | 20304 | 3.43537E-11 | 4.345 |
| *Ccl2* | 20296 | 1.4133E-08 | 4.103 |
| *Cd14* | 12475 | 1.70926E-06 | 3.789 |
| ***Antiviral defense*** | | | |
| *Zbp1* | 58203 | 1.31732E-10 | 3.87 |
| *Rsad2* | 58185 | 5.21588E-07 | 3.15 |
| *Mx1* | 17857 | 1.69343E-23 | 6.661 |
| *Oas1a* | 246730 | 4.51274E-17 | 5.363 |
| *Oas2* | 246728 | 1.56649E-18 | 5.936 |
| *Oas1g* | 23960 | 9.08554E-27 | 6.051 |
| ***Microglial cell activation*** | | | |
| Tlr1 | 21897 | 4.77279E-16 | 5.524 |
| C1qa | 12259 | 5.14353E-06 | 4.698 |
| Itgam | 16409 | 3.21595E-11 | 4.644 |
| Tlr6 | 21899 | 1.2721E-10 | 4.137 |
| Tlr2 | 24088 | 2.54411E-12 | 4.551 |

**B**

| Gene name | ENTREZ ID | Padj. value | Log2FC |
|---|---|---|---|
| ***Cell adhesion*** | | | |
| Pcdghb5 | 93702 | 0.019307769 | -2.515 |
| Lamc1 | 226519 | 0.033045442 | -1.901 |
| Hapln3 | 67666 | 0.005017861 | -3.197 |
| Celsr2 | 53883 | 0.007928181 | -2.024 |
| ***Extracellular matrix*** | | | |
| *Ecm2* | 407800 | 0.027312957 | -2.719 |
| *Eln* | 13717 | 0.04607209 | -3.325 |
| *Dmp1* | 13406 | 0.00918292 | -2.966 |
| *Fbln1* | 14114 | 0.000304742 | -2.65 |
| ***Lipid metabolism*** | | | |
| *Slc27a1* | 26457 | 0.002734674 | -3.727 |
| *Acaa2* | 52538 | 0.017965848 | -1.756 |
| *Msmo1* | 66234 | 0.005265657 | -3.066 |
| *Aacs* | 78894 | 0.046394184 | -1.502 |
| *Fads2* | 56473 | 0.016015922 | -2.626 |
| ***Wnt signaling pathway*** | | | |
| *Yap1* | 22601 | 0.017048647 | -2.16 |
| *Fzd1* | 14362 | 5.3836E-05 | -3.98 |
| *Fzd2* | 57265 | 5.58014E-06 | -4.586 |
| *Wnt7b* | 22422 | 0.000222057 | -2.833 |
| ***Smoothened signaling pathway*** | | | |
| *Evc2* | 68525 | 0.033726815 | -1.934 |
| *Ptch1* | 19206 | 0.036682933 | -2.18 |
| *Tulp3* | 22158 | 0.021707837 | -1.855 |
| *Pax6* | 18508 | 0.002378923 | -2.576 |
| *Gpr37l1* | 171469 | 0.000740359 | -5.326 |
| *Gli1* | 14632 | 0.001120731 | -3.915 |
| *Gli3* | 14634 | 0.027162741 | -2.299 |
| *Gli2* | 14633 | 0.039839325 | -1.732 |
| ***Notch signaling pathway*** | | | |
| *Sox2* | 20674 | 0.00023992 | -4.009 |
| *Dll4* | 54485 | 0.030377179 | -2.442 |
| *Enho* | 69638 | 0.002147326 | -3.159 |
| *Yap1* | 22601 | 0.017048647 | -2.16 |
| *Notch1* | 18128 | 0.03631913 | -1.667 |
| *Hes1* | 15205 | 0.00021596 | -3.237 |
| *Ascl1* | 17172 | 0.033853329 | -2.117 |
| *Hes5* | 15208 | 1.49879E-05 | -5.46 |

**C**

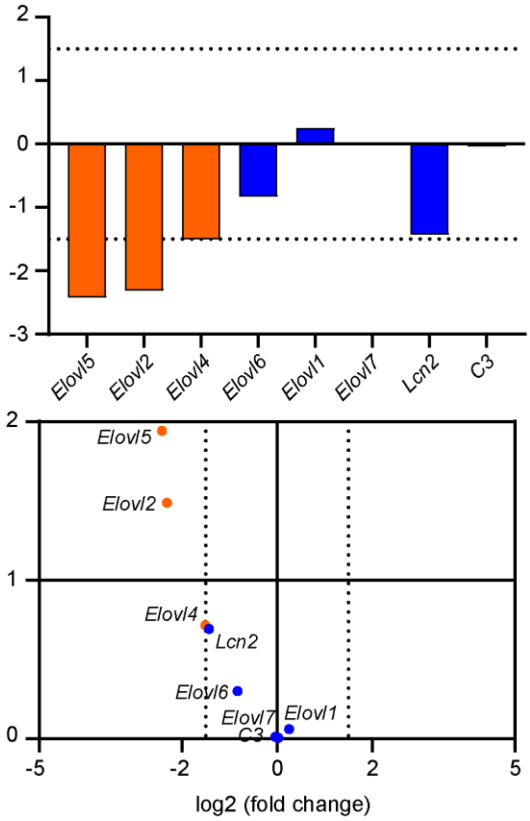

**Figure S7. KEGG pathway analysis of *Srf*-deficient astrocytes.** (**A**) Table showing KEGG pathway and genes upregulated in *Srf*-deficient astrocytes. (**B**) Table showing KEGG pathway and genes downregulated in *Srf*-deficient astrocytes. (**C**) Expression of genes involved in neurotoxicity of reactive astrocytes in *Srf*-deficient astrocytes. Genes with fold change of >2.0 and adjusted p-value < 0.05 were considered significant. Dotted lines indicate cut-off fold change of 1.5.

**Figure S8**

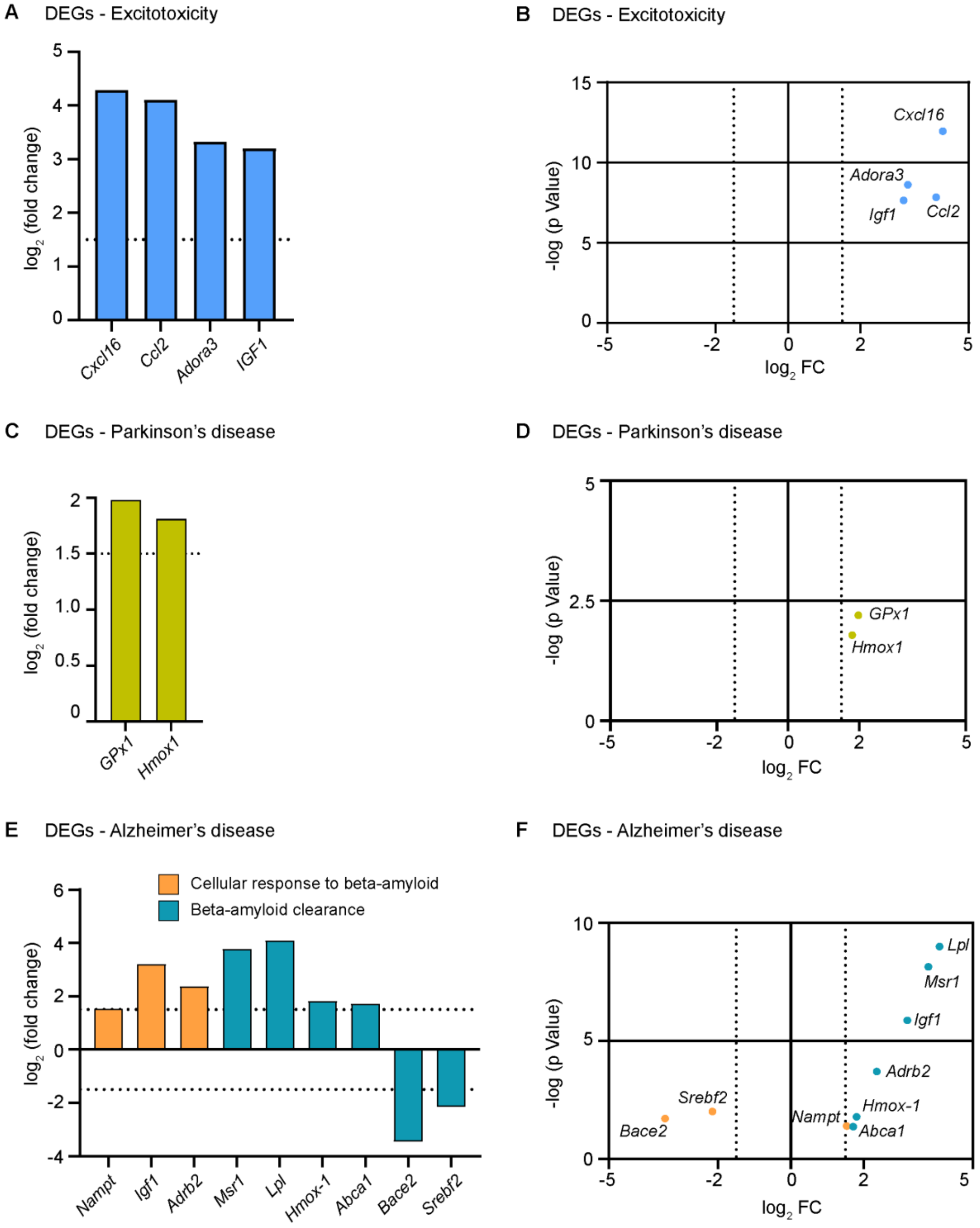

**Figure S8. Differential expression of genes involved in neuroprotection in *Srf* mutant astrocytes. (A)** Bar graph showing the differentially expressed genes (DEGs) from RNA-Seq analysis upregulated in *Srf*-deficient astrocytes that may confer neuroprotection in kainic acid-induced excitotoxicity. **(B)** Scatter plot of DEGs that could protect neurons from kainic acid-induced excitotoxicity. **(C)** Bar graph showing genes upregulated in *Srf*-deficient astrocytes that protect dopaminergic neurons in Parkinson's disease. **(D)** Scatter plot of neuroprotective genes in PD that are upregulated in *Srf*-deficient astrocytes. **(E)** Bar graph showing the upregulated and downregulated genes involved in beta-amyloid cellular response and clearance in *Srf*-deficient astrocytes. **(F)** Scatter plot of DEGs involved in beta-amyloid cellular response and clearance.

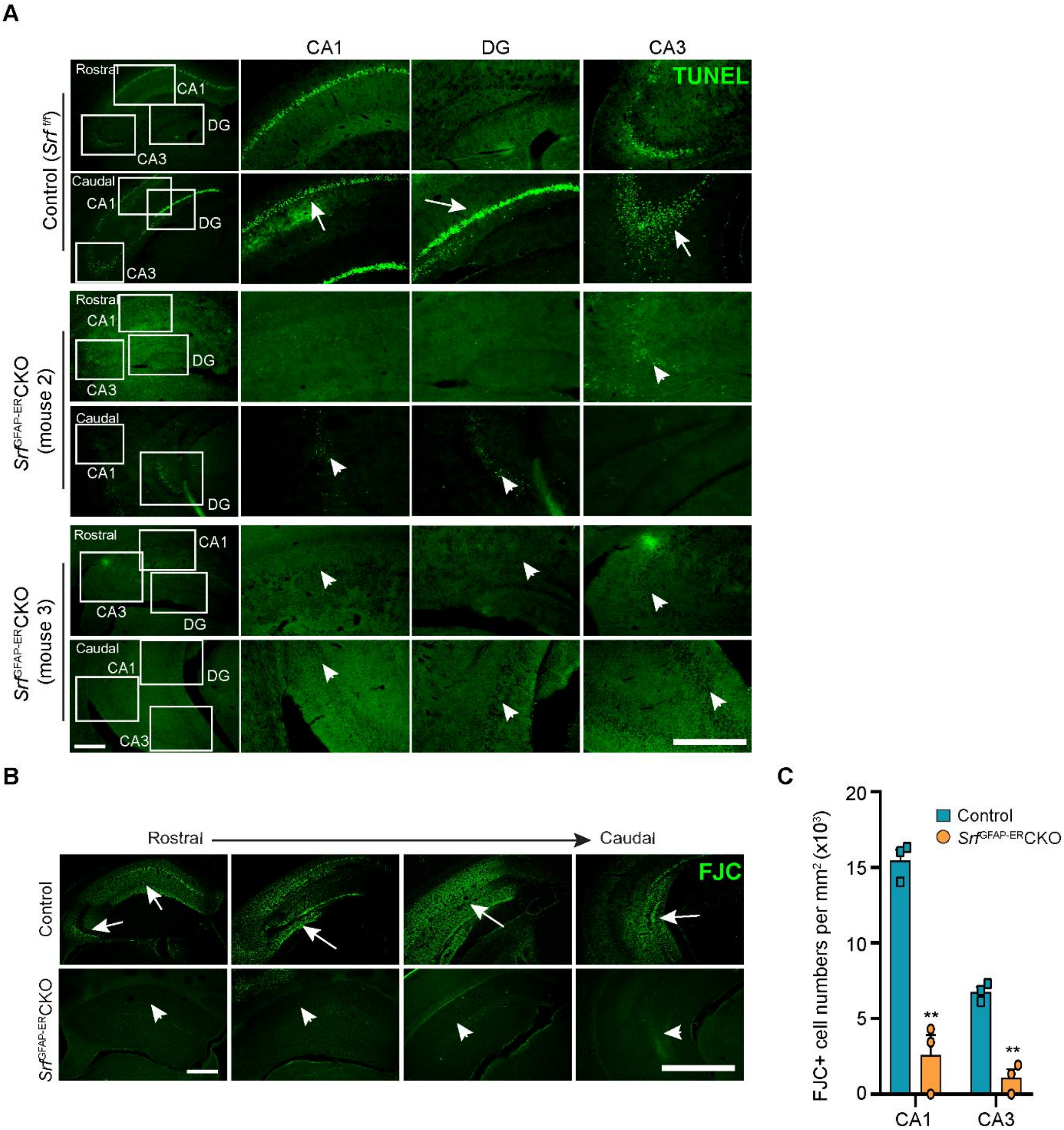

**Figure S9. Neuroprotection from kainic acid-induced excitotoxicity. (A)** Coronal sections showing TUNEL staining of apoptotic cells in the hippocampus of control and *Srf* mutant mice at 4mpi. Representative images from two additional mutant mice are shown. Enlarged images of boxed regions of CA1, CA3 and DG are shown. White arrows and arrowheads show TUNEL+ cells in the control and mutant mice, respectively. **(B)** Coronal sections showing FluoroJade-C (FJC) staining of degenerating neurons and neuronal processes in the hippocampus of control and *Srf* mutant mice at 7-days post-kainic acid administration and at 4 mpi. Shown are the rostral to caudal hippocampal regions. Arrows and arrowheads show the region of degenerating neurons in the control and mutant mice, respectively. **(C)** Quantification of FJC+ cells in the hippocampal CA1 and CA3 regions. n=4 mice. ** $P < 0.01$. Unpaired t-test. Data are represented as mean ± SEM. Scale bar, 200 μm.

**Figure S10**

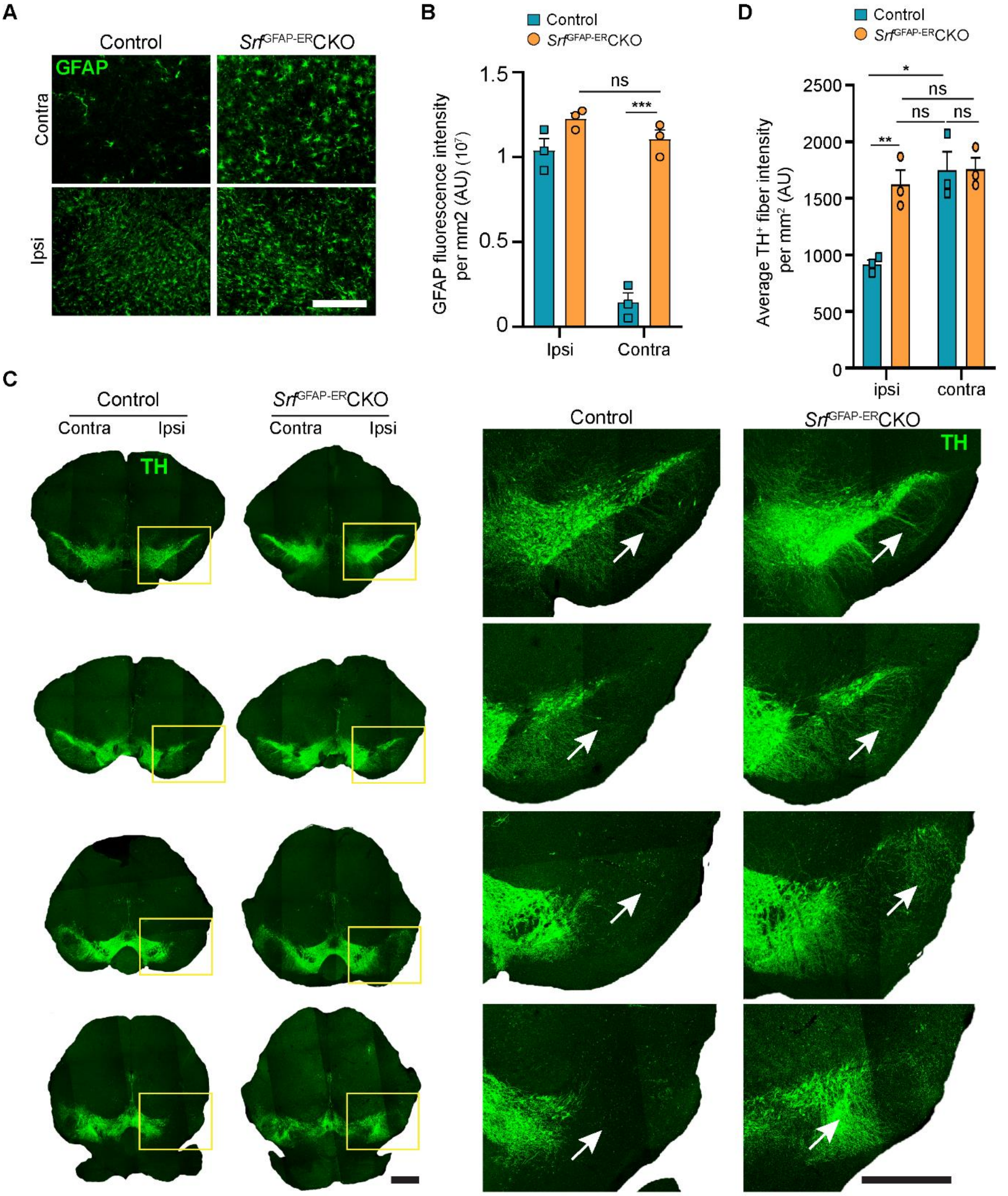

**Figure S10. Neuroprotection in Parkinson's disease mouse model.** **(A)** Tiled confocal images of coronal sections showing immunostaining for tyrosine hydroxylase (TH) in the substantia nigra in control and *Srf*$^{GFAP\text{-}ER}$CKO mice at 10 days post-6-OHDA administration and at 9 mpi. The magnified view of the yellow boxed regions is shown on the right. The white arrows indicate the TH+ fibers from dopaminergic neurons. Scale bar, 500 µm (left panels), 200 µm (right panels). **(B)** Quantification of fluorescence intensity of TH+ fibers in ipsilateral (ipsi) and contralateral (contra) regions of substantia nigra in A. **(C)** Coronal sections showing immunostaining of GFAP in the substantia nigra in control and *Srf*$^{GFAP\text{-}ER}$CKO at 10 days post-6-OHDA administration and at 9 mpi. **(D)** Quantification of GFAP fluorescence in the ipsilateral (ipsi) and contralateral (contra) sides in the substantia nigra in (C). * $P < 0.05$, ** $P < 0.01$, ns, not significant. Unpaired t test. Data are represented as mean ± SEM. AU, arbitrary units.
